# Investigating the function of C-terminal tails of human tubulin isotypes in the motility regulation of cytoplasmic dynein

**DOI:** 10.64898/2026.03.11.711045

**Authors:** Jivesh Garg, Julia Lopes Riberio, Stefan Wallin, Laleh Alisaraie

## Abstract

The intracellular transport system is pivotal for cellular function and integrity, facilitated by cytoskeletal motor proteins such as dynein, which traverse along microtubules (MTs). The heterogeneity of the tubulin isotypes composing MTs introduces functional diversity, potentially affecting cytoskeletal motor proteins’ interactions with the MT. This *in silico* study investigated the influence of amino acid sequence variations in the C-terminal tails (CTTs) of six different *Homo sapiens* tubulin isotypes, TUBB2A, TUBB2B, TUBB2C, TUBB3, TUBB4A, and TUBB5, highly expressed in human brain tumors, and assessed the isotypes’ effect on the binding of motor protein dynein to MT. Among these isotypes, TUBB2A, TUBB2B, and TUBB2C were found to affect conformational motions of the dynein’s microtubule-binding domain (MTBD) and stalk domain. The investigation highlighted the novel role of isotype-specific variations in lateral interactions between tubulin protofilaments (PFs) in determining the proximity of the β-CTT of the adjacent PF to the MTBD, potentially affecting dynein’s motility and suggesting how changes in isotype expression directly influence dynein’s velocity and processivity and contribute to transport defects associated with neurological disorders and cancers. Isolating specific tubulin isotypes experimentally is challenging due to their high sequence similarity and complex interactions with other microtubule-associated proteins. This makes it challenging to distinguish between different tubulin isotypes and their effects, particularly in tissues where multiple isotypes are co-expressed. Additionally, these isotypes are heavily modified *in vivo* by post-translational modifications, which further complicate the isolation of a single, unmodified tubulin isotype. As a result, computational approaches have been necessary in this study for exploring these effects in a controlled, isotype-specific manner.

## 1. Introduction

Microtubules (MTs) are cytoskeletal biopolymers involved in several cellular processes, such as cell division and motility, intracellular transport, and cell shape maintenance (1, 2). MTs comprise α, β-tubulin heterodimers are arranged head-to-tail to form long protofilaments (PFs) that align laterally to shape a hollow cylindrical structure (3). MTs generally consist of thirteen (13) PFs, although variations with a higher (14–16) or lower (9–12) number of PFs have been reported in different species (4). The polarity of an MT arises from the asymmetric arrangement of tubulin subunits, where the slow-growing minus (−) end is characterized by exposed α-tubulin subunits, and the fast-growing plus (+) end by exposed β-tubulin subunits (1). MTs are dynamic structures that continuously undergo cycles of polymerization and depolymerization (5). They also act as platforms along which cytoskeletal molecular motor proteins (e.g., kinesins and dyneins) transport cellular cargo, such as proteins, RNA, and organelles, within cells as they interact with MTs with different binding affinities when in motion. Regulation of MT dynamics is crucial for normal cellular functioning and is primarily achieved through the interaction of MTs with various microtubule-associated proteins (MAPs), including the motor proteins (6). (**Figure 1A**)

**Figure 1:**
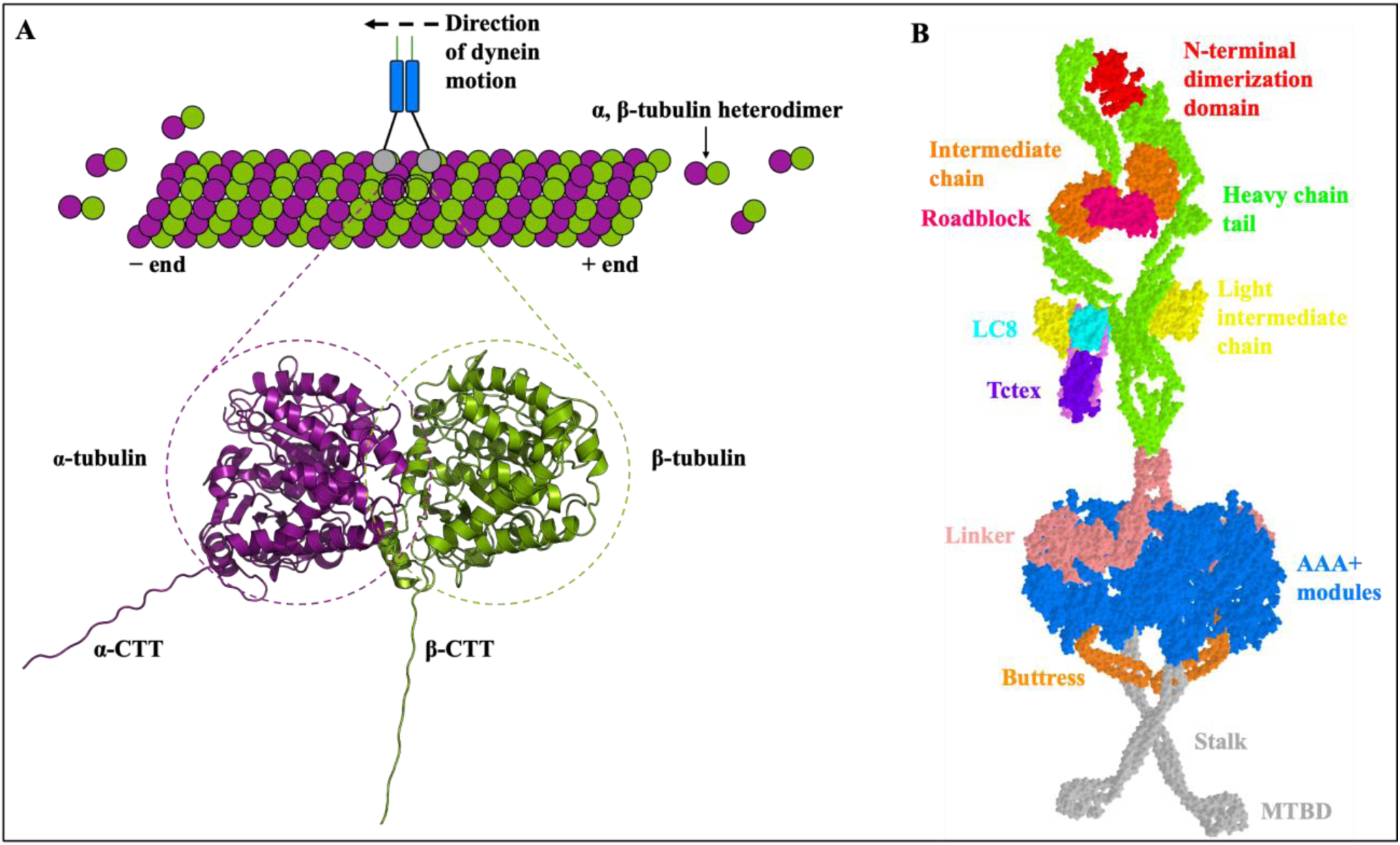
(A) Schematic representation of a microtubule composed of tubulin heterodimers, highlighting the α-subunit (purple) and β-subunit (green), along with their respective C-terminal tails and dynein showing its motion towards the minus (−) end of the MT. (B) Crystal structure of *Homo sapiens* cytoplasmic dynein 1 in its autoinhibited ‘phi’ state conformation, based on the Protein Data Bank entry 5NVU (12).

Dynein is a large (∼1.4 MDa), multi-subunit cytoskeletal motor protein that moves on MTs with the help of its microtubule-binding domain (MTBD) toward MT’s minus (−) end and is involved in a wide range of cellular processes, including cargo transport, cell spindle positioning during cell division, chromosome segregation, and organization of cilia and flagella (7). Dyneins are members of the AAA^+^ superfamily of ATPase enzymes that convert the energy released from ATP hydrolysis into mechanical movement, enabling them to move along MTs (8). One of the two cytoplasmic dyneins, dynein 1 (hereafter referred to as dynein), comprises two identical heavy chains with a hexameric head (motor domain), linker, stalk, and MTBD, two light intermediate chains, four intermediate chains, and six light chains (two copies of each LC8, Roadblock, and Tctex) (9). The motor domain of dynein comprises six AAA^+^ (AAA1–AAA6) modules that form a ring with a diameter of approximately 13 nm, converting the chemical energy released from ATP hydrolysis into mechanical motion (7). The four AAA^+^ modules (AAA1–AAA4) can bind to a nucleotide, and AAA1 is the primary ATP-binding domain (9). The stalk is a long (∼15 nm) coiled coil that consists of two antiparallel α-helices extending from the hexameric head, with its terminal connected to MTBD, which enables dynein to move on MTs (10). The MTBD interacts with microtubules through different binding modes, known as the α-registry and β-registry, associated with a strong (high-affinity) and weak (low-affinity) binding states of the MTBD to the MT, respectively (11). (**Figure 1B**)

The diversity and regulation of MTs are further enhanced by multiple tubulin isotypes, each encoded by distinct genes. For instance, *Homo sapiens* consists of at least nine α-tubulin and ten β-tubulin isotypes (13), which exhibit varied expression profiles and tissue-specific expression patterns (14). While these isotypes are highly conserved overall (>90% sequence identity), their primary differences lie in the length, composition, and arrangement of the amino acids in their C-terminal tails (CTTs). These different tubulin isotypes and various post-translational modifications (PTMs) govern microtubule (MT) characteristics, stability, and dynamics, a notion known as the “tubulin code” (5). Some isotypes, such as αIA, βI, and βV, are ubiquitously expressed, whereas others are limited to specific cell types or tissues. For example, βIIA, βIIB, βIII, and βIVA are predominantly found in the brain, while βVI is mainly present in erythroid cells and platelets (14). Their expression levels and distributions can also vary throughout development (15) and the cell cycle (16) to fulfill specific cellular requirements. βIII-tubulin is abundantly expressed in neurons and the embryonic nervous system, with its expression typically decreasing in non-neuronal cells as they undergo differentiation (15, 17). αIA-tubulin is highly expressed in post-mitotic neurons but declines in later developmental stages (15), while αIVA-tubulin is prominent in platelets, especially during late stages of megakaryocyte differentiation (18). During the cell cycle, Class IV β-tubulin isotypes (IVA and IVB) show elevated transcript and protein levels in dividing cells compared to resting cells, suggesting their specialized role in mitotic spindle dynamics (16). Moreover, the expression of tubulin isotypes is altered in various diseases, particularly in cancers and in response to drug treatments (14). Notably, βIII-tubulin is frequently overexpressed in advanced and metastatic tumors, where its elevated levels are associated with resistance to microtubule-targeting agents (MTAs) such as taxanes and vinca alkaloids, and are often linked to poor clinical outcomes (19). Tubulin isotypes also influence various MT properties, including dynamic instability, polymerization rates, catastrophe frequencies, and growth/shrinkage rates, which can indirectly affect how motor proteins interact with the MT lattice. For instance, αIC/βIIA MTs exhibit faster growth and lower catastrophe frequencies compared to αIA/βIIA MTs (20). Conversely, βIII-tubulin has been associated with more dynamic MTs and higher catastrophe frequencies (17). Isotypes also regulate MT structure, including PF number and lattice architecture. For example, *Homo sapiens* αIB/βIII isotypes predominantly form 13-PF MTs, while αIB/βIIB isotypes mainly form 14-PF MTs (21). Microtubules composed of αIVA/βIIA exhibit longitudinal contraction compared to those with αIA/βIIA and αIC/βIIA (22). Furthermore, MTs can display sectioned structures where isotype composition and PF numbers vary along their length, thereby affecting interactions with MAPs, including motor proteins (23).

The dynamic interaction between dynein and the MT track is paramount for its function, and this interaction can be modulated by the intrinsic properties of the microtubule itself, notably its isotype composition and the C-terminal tails (CTTs) of α- and β-tubulin (aka E-hook). While the CTTs are not the primary binding sites for dynein, they nonetheless significantly influence dynein’s interaction with MTs, primarily through the electrostatic interactions, since these CTTs have a high proportion of negatively-charged amino acids (namely, glutamates and aspartates) (24). Moreover, tubulin CTTs can undergo multiple post-translational modifications (PTMs), such as polyglutamylation, polyglycylation, and tyrosination/detyrosination, and display sequence variability among tubulin isotypes, which can affect MT stability and function, including the binding affinity of motor proteins to MTs (25). Recent studies have suggested that the tubulin CTTs regulate dynein’s activity, either directly through electrostatic interactions with the MTBD (26, 27) or indirectly through MAPs, including MAP9 (28) and MAP4 (29). Several lines of evidence, both computational and experimental, support a direct role of the β-CTT in regulating the binding and processivity of dynein’s MTBD to the MT segments (30, 31). A molecular dynamics (MD) simulation study of the α, β-tubulin heterodimer bound to the MTBD suggested that a number of amino acids from both β-CTT (*Sus scrofa*) and MTBD (*Dictyostelium discoideum*) affect MTBD binding at its α/β registries through salt bridges, electrostatic interactions, and hydrogen bonds (26). E-hook was shown to interact with MTBD helices H1 and H6, resulting in the shift of helix H1 to a nearly parallel position relative to heterodimer tubulin that can lead to conformational change required for the MTBD switching between low and high binding affinities (26). Another MD study indicated that E-hooks (*Bos taurus*) guide MTBD (*Mus musculus*) towards the binding position on α, β-tubulin heterodimer via direct (viasalt bridges) and long-range electrostatic interactions (24). Using bead motility assays, Wang and Sheetz found that the removal of CTTs from MTs (porcine brain) decreased dynein (chicken embryo brain) binding to MTs by three times, its diffusion by twenty times, and reduced run length of the dynein by fourfold, pointing to the involvement of the E-hook in maintaining contacts with dynein (30). Additionally, the C-terminal region of tubulin (*Bos taurus*) has been implicated in stimulating cytoplasmic dynein’s (*Bos taurus*) ATPase activity, as removal of CTTs from MTs led to a five- to sixfold decrease in the microtubule-activated ATPase activity of the dynein (31). In another study, removal of the E-hooks (*Bos taurus*) caused a two-fold reduction in the run length of yeast dynein (*Saccharomyces cerevisiae*) (32). Sirajuddin *et al.* (27) reported that yeast (*Saccharomyces cerevisiae*) dynein exhibited comparable velocity and processivity on yeast MTs, porcine MTs, and recombinant MTs (TUBA1A/TUBB2A) consisting of yeast tubulin core fused with *Homo sapiens* CTTs. Furthermore, removal of both the α- and β-CTTs did not affect the velocity of yeast dynein but reduced dynein processivity by 50%. The β-CTT restored this loss in processivity, whereas the α-CTT had no such effect. Among the human tubulin CTTs tested (TUBB1, TUBB2A, TUBB3, TUBB4A, TUBB6, TUBB7, and TUBB8), only the TUBB3 CTT increased the run length of yeast dynein by approximately 1.5-fold, while velocity remained unaffected across all isotypes (27). These results suggest that CTTs enhance yeast dynein motility, but are not strictly required, as yeast dynein moves almost as well on CTT-truncated MTs as it does on untreated MTs (27), likely because yeast dynein is inherently processive without requiring any other cofactors or adaptor proteins (33). However, purified mammalian dynein is largely inactive, adopting a “phi-particle” conformation incapable of robust, processive MT binding and ATP hydrolysis. Instead, mammalian dynein requires the simultaneous binding of dynactin and specific cargo adaptor proteins (such as Bicaudal D cargo adaptor 2 (BicD2), Spindly, or Hook3) to become highly processive and move over long distances (12). Therefore, mammalian dynein may interact with the E-hooks differently, as shown by McKenney *et al.* (34), that removal of tubulin CTTs (*Sus scrofa*) by subtilisin significantly reduced human recombinant DDB (Dynein−Dynactin−BicD2) construct binding and nearly abolished its processive movement, in contrast to its robust motility on untreated MTs (34). This indicates that, unlike yeast dynein, mammalian dynein motility depends critically on the presence of the tubulin CTTs.

Despite increasing research highlighting the role of tubulin CTTs in regulating dynein activity, the detailed mechanisms, especially in *Homo sapiens*, remain elusive. Most previous studies have investigated the effect of tubulin CTTs on dynein in species such as *Saccharomyces cerevisiae* (27, 32), *Mus musculus* (24), *Bos taurus* (24, 31), and *Sus scrofa* (26, 30, 34). Many of these studies also combined MTs from one species with MTBDs (or dynein) from another (24, 26, 30, 32, 34). Although these species are evolutionarily related to *Homo sapiens*, their tubulin CTT sequences differ, highlighting the need for species-specific investigations. Furthermore, the influence of different tubulin isotypes on dynein remains poorly understood. While a wide variety of kinesin family motors exist to drive transport toward the MT plus-end, there is only one major cytoplasmic dynein responsible for all minus-end-directed transport, and it remains unclear how dynein’s motility is regulated across different tissues, a regulation that depends on the specific tubulin isotypes expressed in each tissue. Key questions remain unanswered, especially regarding how variations in the CTT’s sequences among tubulin isotypes influence dynein binding to the MTs, the resultant configurational changes in the MTBD, and their impacts on dynein’s velocity and processivity.

Isolating specific tubulin isotypes experimentally is challenging due to their high sequence similarity and complex interactions with other MAPs. This makes it challenging to distinguish between different tubulin isotypes and their effects, particularly in tissues where multiple isotypes are co-expressed. Additionally, these isotypes are heavily modified *in vivo* by PTMs, which further makes it difficult to isolate a single, unmodified tubulin isotype. As a result, computational approaches are necessary to explore these effects in a controlled, isotype-specific manner.

Utilizing Molecular Dynamics (MD) simulations, Principal Component Analysis (PCA), and Dynamical Cross-Correlation (DCC) analysis, this research emphasizes β-tubulin isotypes, specifically *Homo sapiens* TUBB2A, TUBB2B, TUBB2C (TUBB4B), TUBB3, TUBB4A, and TUBB5 (TUBB), and their E-hooks, exploring their roles in dynein binding to the MT. This study provides structural evidence that E-hooks can have different effects on the MT-MTBD interactions, even in closely related species (such as *Homo sapiens*, *Bos taurus*, and *Sus scrofa*), due to differences in their amino acid sequences at the CTT region. Additionally, this work examines the interactions between the β-CTT of the adjacent PF and the MTBD in different isotypes, as well as the effect of these interactions on the binding profile of the MTBD. It explores the correlation between the interdimer angle between the laterally binding PFs (PF_1_ and PF_2_) and the involvement of the MTBD-bound α, β-tubulin heterodimer in PF_1_ and the β-CTT of the adjacent heterodimer in PF_2_. The results suggest a novel role and functional significance of lateral interactions between the PFs, which can affect MT-involved processes. By investigating conformational changes in the involved domains, this study shows how variations in the isotype-specific amino acid sequence affect lateral interactions of MT protofilaments (e.g., as in PF_1_ and PF_2_) and, consequently, can influence the conformational changes of MTBD required for dynein’s motility.

## 2. Materials and Methods

### 2.1. Preparation of the initial tubulin structures

The initial tubulin protein structures used in this study were retrieved from the Protein Data Bank entry 5JCO (35), featuring a cryo-electron microscopic solved structure of *Homo sapiens* post-translationally unmodified αIA(TUBA1A)/βIII(TUBB3) MTs (*Homo sapiens*, UniProt code Q71U36 (36) (α-tubulin) and Q13509 (37) (β-tubulin)) (hereafter referred to as wild type) with a resolution of 4.00 Å. The initial tubulin structure (5JCO) (35) encompassed 12 subunits (six α-tubulin and six β-tubulin) with the nucleotides (guanosine-5’-triphosphate (GTP) and phosphomethylphosphonic acid guanylate ester (GMPCPP) in the α- and β-tubulin subunit, respectively). However, in this study, the number of tubulin subunits was truncated to either a dimer, composed of a subunit of each α- and β-tubulin, or a tetramer comprising two subunits of each α- and β-tubulin, representing PF_1_ and PF_2_. (**Figure 2 and Figure 3**)

**Figure 2:**
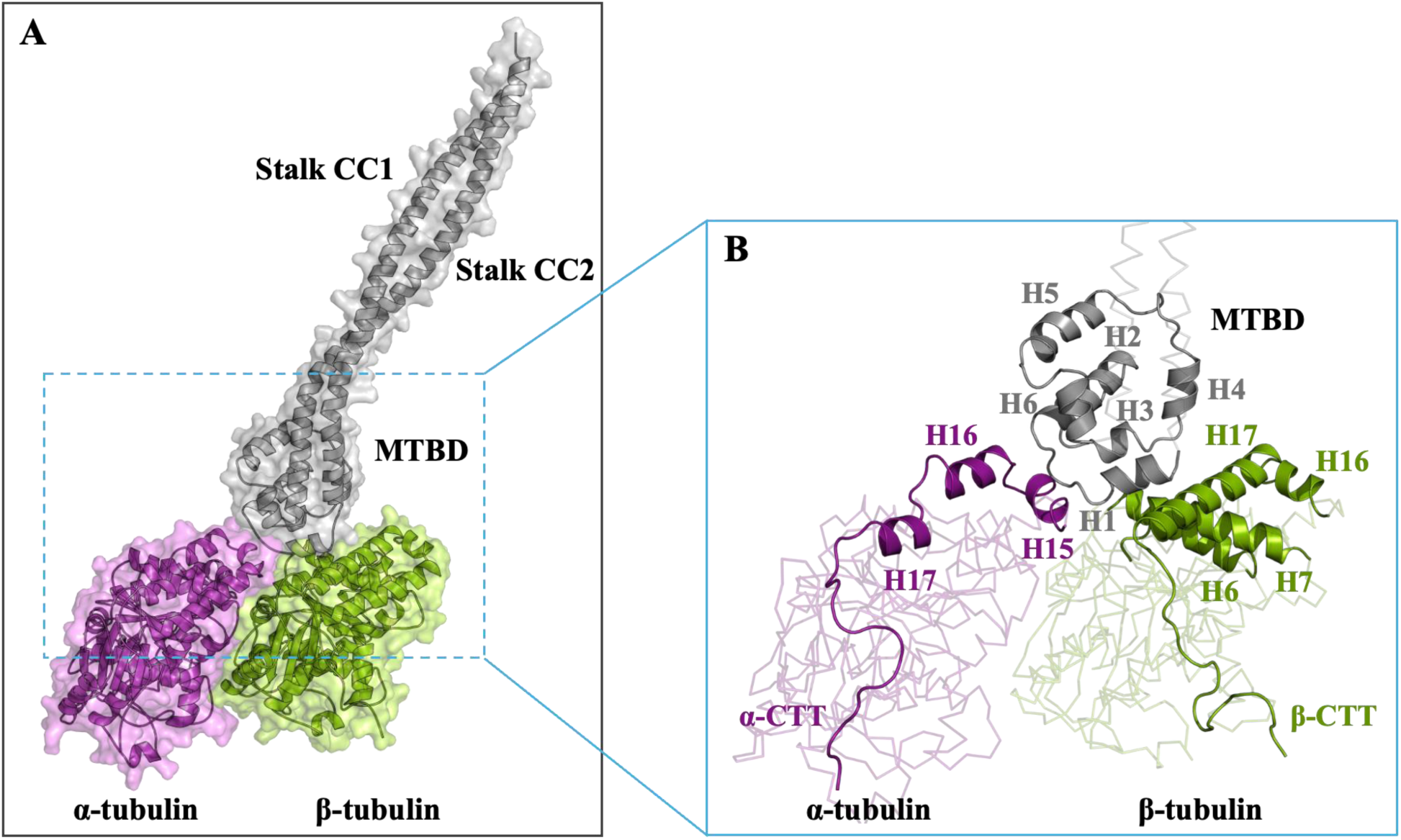
(A) Structural schematic of the α, β-tubulin heterodimer, showing α-tubulin (purple), β-tubulin (green), and the dynein microtubule-binding domain (MTBD) with its stalk domain (gray). (B) Helices of α-tubulin (H15, H16, H17) and β-tubulin (H6, H7, H16, H17) that form the MTBD-binding interface, alongside the six helices of the MTBD (H1–H6).

**Figure 3:**
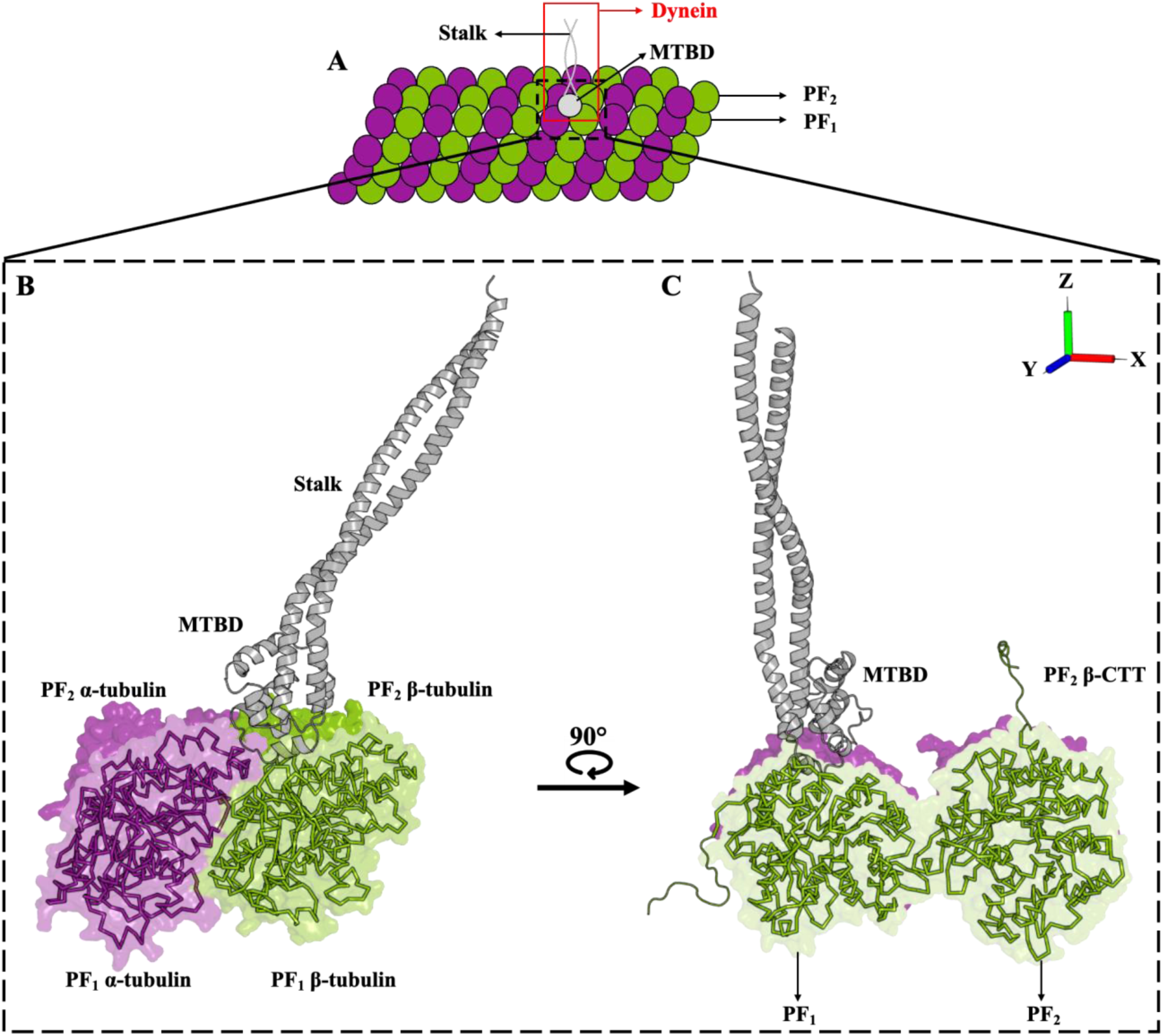
(A) Cartoon representation of a microtubule indicating protofilaments PF_1_ and PF_2_ and dynein’s MTBD and stalk domains. (B) and (C) Schematic of the tubulin tetramer consisting of two α-tubulins (purple) and β-tubulins (green) arranged in two protofilaments (PF_1_ and PF_2_) with dynein’s MTBD and stalk domain (gray) bound to the α, β-heterodimer of PF_1_.

Notably, amino acids ranging from 38–46 and 438–451 (i.e., α-CTT) within the α-tubulin subunit and amino acids ranging from 427–450 (i.e., β-CTT) within the β-tubulin subunit were not characterized (35) due to their high flexibility, known as an intrinsically disordered region (IDR) (38). Thus, in this work, the CTTs and any other uncharacterized residues were first modeled and merged with the tubulin built up in 5JCO (35).

The α1A tubulin (TUBA1A) in 5JCO comprises 451 amino acids, characterized by 17 α-helices (H1–H17) and 12 β-strands (E1–E12). The βIII tubulin isotype (TUBB3), sourced from the Protein Data Bank (5JCO), comprises 450 amino acids, featuring 17 α-helices (H1–H17) and 11 β-strands (E1–E11). The proteins’ secondary structure assignments were determined using the Protein Secondary Structure Visualiser (ProS^2^Vi) software package, which functions based on the Dictionary of Secondary Structure in Proteins (DSSP) algorithm (39). (**Figure S1A–B**)

The β-tubulin amino acid sequence information of *Homo sapiens* tubulin isotypes, namely TUBB2A, TUBB2B, TUBB2C (TUBB4B), TUBB4A, and TUBB5 (TUBB), was sourced from UniProt entries Q13885 (40), Q9BVA1 (40), P68371 (41), P04350 (42), and P07437(43), respectively. The isotypes’ tertiary structures were modeled according to tubulin heterodimer units and their arrangements in 5JCO. These models were then used as the starting protein isotype structures for molecular dynamics (MD) simulations.

Sequence alignment of the wild-type β-tubulin (TUBB3) and β-tubulin isotypes (TUBB2A, TUBB2B, TUBB2C, TUBB4A, and TUBB5) was conducted using the Clustal Omega algorithm available through UniProt (40, 44). TUBB2A shows 91.46% sequence identity (sharing 407 of the 445 amino acids), TUBB2B has 91.69% identity (i.e., 408 out of the 445 amino acids), TUBB2C has 92.81% identity (413 out of 445 amino acids), TUBB4A has 92.12% identity (i.e., 409 out of the 444 amino acids) and TUBB5 has 92.57% sequence identity (sharing 411 of the 444 amino acids) with the wild type (TUBB3). Notable differences are in the isotypes’ CTTs. The CTTs of both TUBB2A and TUBB2B isotypes exhibit 47.37% sequence identity (9 identical amino acids out of 19), TUBB2C CTT shows 68.42% identity (with 13 amino acids identical out of 19), TUBB4A CTT has 66.67% identity (each with 12 amino acids identical out of 18), and TUBB5 CTT has 61.11% sequence identity (11 identical amino acids out of 18) with the TUBB3 CTT. (**Figure S2**)

The structures of the MTBD and stalk domain of dynein were retrieved from the Protein Data Bank with the 5AYH code (*Mus musculus,* UniProt code Q9JHU4) (10). This structure was used as a template for homology modeling the MTBD and the stalk domain according to the *Homo sapiens* sequence information (UniProt entry Q14204) due to the lack of an experimentally characterized structure of *Homo sapiens* MTBD attached to the stalk. Both *Mus musculus* and *Homo sapiens* MTBD and the stalk domain showed 100% sequence identity in their 254 amino acids, making it a suitable template for modeling the *Homo sapiens* MTBD and the stalk domain. (**Figure S3**)

The resultant *Homo sapiens* MTBD and the stalk domain were aligned to the experimentally determined bound position of the MTBD (*Dictyostelium discoideum,* UniProt code P34036) to α, β-tubulin heterodimer (*Sus scrofa*, TUBA1A/TUBB) according to the pseudo-atomic model of the MTBD-tubulin heterodimer complex in 3J6P (45), serving as the template structure for modeling the MTBD bound to the α, β-heterodimer in PF_1_. (**Figure 2 and Figure 3**)

The MTBD and stalk domains encompass a total of 254 residues (Asp3221–Arg3474) with MTBD spanning amino acids Lys3301–Ala3385 alongside a stalk subunit comprising two coiled coils (CC), namely, CC1 (amino acids Arg3223–Lys3297) and CC2 (amino acids Cys3389–Val3472). Six helices were identified within the MTBD, denoted as H1 (Lys3301–Arg3308), H2 (Ala3315–Leu3327), H3 (Trp3335–Ile3341), H4 (Phe3347–Val3352), H5 (Asp3361–Tyr3371), and H6 (Tyr3379–Ala3385), according to the ProS^2^Vi software (39). (**Figure 2B and Figure S1C**)

### 2.2. Preparation of the different tubulin molecular systems

Twelve distinct tubulin molecular systems (S1–S12) were prepared for separate MD simulations based on four key factors: (i) *Homo sapiens* β-tubulin isotype sequence information of five (5) different β-tubulin isotypes, TUBB2A, TUBB2B, TUBB2C, TUBB4A, and TUBB5, and its wild-type TUBB3. (ii) number of the α- and β-tubulin subunits, either as a dimer in PF_1_ or a tetramer comprising PF_1_ and PF_2_, where the MTBD-stalk domain bound to the α, β-tubulin heterodimer is PF_1_, and the tetramer includes PF_2_ adjacent to PF_1_. (iii) the implemented force field, GROMOS 54A7 vs CHARMM36. Chemistry at HARvard Macromolecular Mechanics (CHARMM36) is an all-atom force field, explicitly treating and parameterizing every atom, while GROningen MOlecular Simulation (GROMOS 54A7) is a united-atom force field (46), grouping non-polar hydrogen atoms with carbon atoms and assigning parameters to the resultant CH, CH_2_, or CH_3_ ‘pseudo-atoms.’ United-atom force fields significantly reduce the number of atoms for which non-bonded interactions need to be calculated, thus decreasing the computational expense by up to an order of magnitude (47). (iv) the presence or absence of the nucleotides in the α, β-tubulin heterodimer, specifically guanosine-5’-triphosphate (GTP) in the α-subunit and phosphomethylphosphonic acid guanylate ester (GMPCPP) in the β-tubulin subunit. These different molecular systems were prepared to investigate the influence of tubulin isotype-specific CTTs on dynein (i.e., MTBD and stalk domains) and the impact of neighboring β-tubulin’s CTTs on MTBD binding. (**Table 1**)

**Table 1:** Molecular systems (S1–S12), β-tubulin isotypes, and force fields in this study.

| Molecular system | β-tubulin isotype | Molecular systems Composition | Force field | Presence of nucleotide | Convergence time |
| --- | --- | --- | --- | --- | --- |
| S1 | TUBB2B | Dimer | GROMOS 54A7 | Yes | 95 ns |
| S2 | TUBB2B | Dimer | CHARMM36 | Yes | 125 ns |
| S3 | TUBB3 | Dimer | GROMOS 54A7 | Yes | 80 ns |
| S4 | TUBB3 | Dimer | CHARMM36 | Yes | 100 ns |
| S5 | TUBB5 | Dimer | GROMOS 54A7 | Yes | 60 ns |
| S6 | TUBB5 | Dimer | CHARMM36 | Yes | 60 ns |
| S7 | TUBB2A | Tetramer | GROMOS 54A7 | No | 150 ns |
| S8 | TUBB2B | Tetramer | GROMOS 54A7 | No | 140 ns |
| S9 | TUBB2C | Tetramer | GROMOS 54A7 | No | 110 ns |
| S10 | TUBB3 | Tetramer | GROMOS 54A7 | No | 260 ns |
| S11 | TUBB4A | Tetramer | GROMOS 54A7 | No | 80 ns |
| S12 | TUBB5 | Tetramer | GROMOS 54A7 | No | 140 ns |

### 2.3. Molecular dynamics (MD) simulations setup

MD simulations were conducted to investigate dynamics and conformational changes in the tubulin subunits, MTBD, and stalk. These simulations were executed using GROMACS (48, 49) version 2022.2. GROMACS works on the principles of classical mechanics to model the motion of a group of atoms. Essentially, the program solves the Newtonian equation of motion, encompassing all atoms constituting the molecular system (50). By virtue of this computational framework, the software offers a robust platform for simulating and analyzing the intricate behaviors of molecular assemblies. To comprehensively evaluate the impact of β-CTT on dynein, two different force fields, namely GROMOS 54A7 (51) and CHARMM36 (52), were used. The GROMOS 54A7 force field was previously employed to study the effect of β-CTT on alterations in the dynein binding mode (26).

The α, β-tubulin heterodimers (S1–S6) also contained nucleotides, specifically guanosine-5’-triphosphate (GTP), in the α-subunit and phosphomethylphosphonic acid guanylate ester (GMPCPP) in the β-subunit. These nucleotides were assigned a −4 charge, accounting for the deprotonation of the nucleotides under physiological conditions. For the 54A7 force field, the nucleotides’ topology was generated using the Automatic Topology Builder (ATB) version 3.0 (53, 54). For the CHARMM force field, the ligand’s topology was generated through the CHARMM General Force Field (CGenFF) program (55), version 2.5.

Each molecular systems (i.e., S1–S12), consisting of the protein and nucleotides, were immersed in a separate cubic simulation box for which the simple point charge (SPC) water model (56) was used, and sodium or chloride ions were added depending on the resulting net charge of each system to neutralize the net charge of the system. Each tubulin molecular system (S1–S12) was centered at a distance of 1.0 nm from the edge of the box. (**Table S1**)

Energy minimization was performed in multiple steps using the steepest descent algorithm (57) to eliminate any atomic clashes. Each system underwent 50,000 steps of energy minimization, ensuring that the maximum force tolerance remained below 1000 kJ mol^-1^ nm^-1^. (**Table S2**)

Periodic Boundary Conditions (PBC) were applied to remove the boundary effects (58). All the molecular systems (S1–S12) underwent a two-stage equilibration process of temperature and pressure involving NVT (constant number of particles, volume, and temperature), followed by NPT (constant number of particles, pressure, and temperature) ensembles. For the NVT ensemble, the V-rescale temperature-coupling method (59) was used. This involved a constant coupling for 1 ns with a step size of 0.002 ps at 300 K. During the NPT phase, the C-rescale pressure-coupling technique (60) was enabled with a consistent coupling duration of 10 ns at 300 K. The short range (i.e., up to 1.4 nm and 1.2 nm in GROMOS 54A7 and CHARMM36, respectively) electrostatic forces were calculated directly while for long-range electrostatic interactions (beyond 1.4 nm with a 0.12 nm Fourier grid spacing in GROMOS 54A7 and 1.2 nm with a 0.16 nm Fourier grid spacing in CHARMM36), Particle Mesh Ewald (PME) method was employed (61, 62). For the van der Waals (vdW) interactions, cutoffs of 1.4 nm and 1.2 nm were used for GROMOS 54A7 and CHARMM36, respectively. The LINear Constraint Solver (LINCS) algorithm (63) was applied to restrain bonds. Notably, position restraints were implemented for all bonds between the hydrogens and heavy atoms for CHARMM36. In contrast, for GROMOS 54A7, all chemical bonds (i.e., between heavy atoms as well) were constrained. Random seeds were selected to ensure integrity. The compressibility was set to 4.5 × 10^−5^ bar, ultimately converging to a steady pressure of 1.0 bar. An MD (leap-frog) integrator was employed during the data collection phase of the MD simulations (58). The conformation obtained after the 10 ns NPT equilibration was used as the starting conformation for the MD simulations. The calculations were conducted over a duration of 1.0 μs for each molecular system using high-performance computing clusters provided by the Digital Research Alliance of Canada and ACENET. The integration time step was fixed at 0.002 ps. This allowed for a detailed examination of system dynamics and behavior over an extended temporal scale.

To assess the convergence of the S1 to S12 systems, the root mean square deviation (RMSD) of the backbone atoms was calculated throughout the trajectory. The tubulin dimers (S1–S6) converged earlier than the tubulin tetramers (S7–S12) due to the molecular systems’ smaller size (1.18 million atoms in dimers vs 1.38 million atoms in tetramers). All the systems converged within 60 ns to 260 ns and were allowed for an additional (up to) 1 µs after convergence; however, the analyses were conducted after convergence. (**Table 1 and Figure S4**)

### 2.4. Principal Component Analysis (PCA) and Dynamic Cross-Correlation (DCC) analysis

To characterize the global collective motions of the TUBB2A, TUBB2B, and TUBB2C tubulin tetramers, PCA, also known as Essential Dynamics (ED), was performed. PCA is a statistical method widely used in the analysis of MD simulations to reduce the dimensionality of large datasets while retaining the most significant dynamical features (64). It achieves this by transforming atomic positional fluctuations into a new orthogonal basis set, where the first few principal components (PCs) capture the majority of the system’s motion (65). For each tetramer (TUBB2A, TUBB2B, and TUBB2C), a covariance matrix of atomic positional fluctuations was computed from a 1 μs MD trajectory using the backbone atoms (N, Cα, C) after least-squares fitting to the last frame of NPT equilibration reference structure. The resulting covariance matrix was then diagonalized to obtain a set of eigenvectors and corresponding eigenvalues, with the eigenvectors ordered according to the magnitude of their eigenvalues (64). The eigenvectors associated with the largest eigenvalues represent the principal modes of motion, contributing most significantly to the protein’s conformational dynamics (66).

In parallel, Dynamic Cross-Correlation (DCC) analysis was performed to quantify the correlated and anti-correlated motions among the two protofilaments (PF_1_ and PF_2_), MTBD, and the stalk domain. The DCC matrix was calculated using Cα atoms by transforming the covariance matrix obtained from PCA into a cross-correlation matrix through a modified Python script (67) adapted for this system. The cross-correlation coefficient between atoms i and j was calculated as follows (68). (**Eq. 1**)

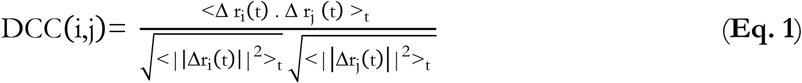

where r_i_(t) denotes the position vector of the i^th^ atom at time t, <>_t_ the time average over the trajectory, and Δr_i_(t)= r_i_(t)-(<r_i_(t)>)_t_ the deviation at time t of atom i from its average position, <r_i_(t)>_t_.

The DCC(i,j) can take values from −1 to 1, where positive values indicate correlated motion (atoms move in the same direction), negative values reflect anti-correlated motion (atoms move in opposite directions), and values near zero suggest uncorrelated motion.

### 2.5. Calculation of the interdimer angle (θ) between laterally interacting protofilaments PF_1_ and PF_2_ in tubulin tetramers

The conformational differences between the wild-type (TUBB3) tetramer and tubulin isotypes (TUBB2A, TUBB2B, TUBB2C, TUBB4A, and TUBB5) at the lateral interface were analyzed in terms of the interdimer angle between the adjacent PFs (PF_1_ and PF_2_) using an in-house Python script utilizing the MDAnalysis package (69).

The interdimer angle was defined by fitting two planes: Plane_1_ through α, β-tubulin subunits in PF_1_ and Plane_2_ through the α, β-tubulin subunits in PF_2_, and computing the angle between their normal vectors, (i.e., 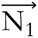 for Plane_1_ and 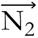 for Plane_2_), building up based on Peng *et al* (70). The singular value decomposition (SVD) algorithm was used to obtain the optimal plane and its normal vector (71, 72). SVD was chosen over other conventional plane-fitting methods, such as eigenvalue decomposition using principal component analysis (PCA) or ordinary least squares (OLS), because it is faster, precise, robust, and numerically stable (73, 74). This is because SVD minimizes the orthogonal distances from all atoms to the fitted plane, making it more robust and unbiased than OLS, which is sensitive to orientation and minimizes distances only along a chosen axis (e.g., the z-axis), making the result dependent on the coordinate system and highly susceptible to noise in that specific direction (75, 76). Similarly, SVD offers higher numerical stability than PCA because it avoids explicit computation of the covariance matrix, which can introduce rounding errors in large coordinate datasets (73). Additionally, the SVD algorithm is also implemented in the PyMOL script (orientation.py (77)) for the calculation of the plane orientation and angle between different domains of a biomolecule, reinforcing its suitability for biomolecular structural analysis. To compute each plane and its normal vectors, the atomic coordinates of α, β-tubulin from PF_1_ (for 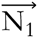) and PF_2_ (for 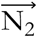) were extracted, centered around the centroid, and subjected to the singular value decomposition. The vector corresponding to the smallest singular value, representing the axis of least variance, was selected as the normal vector of the best-fit plane (71). (**Figure 4**)

**Figure 4:**
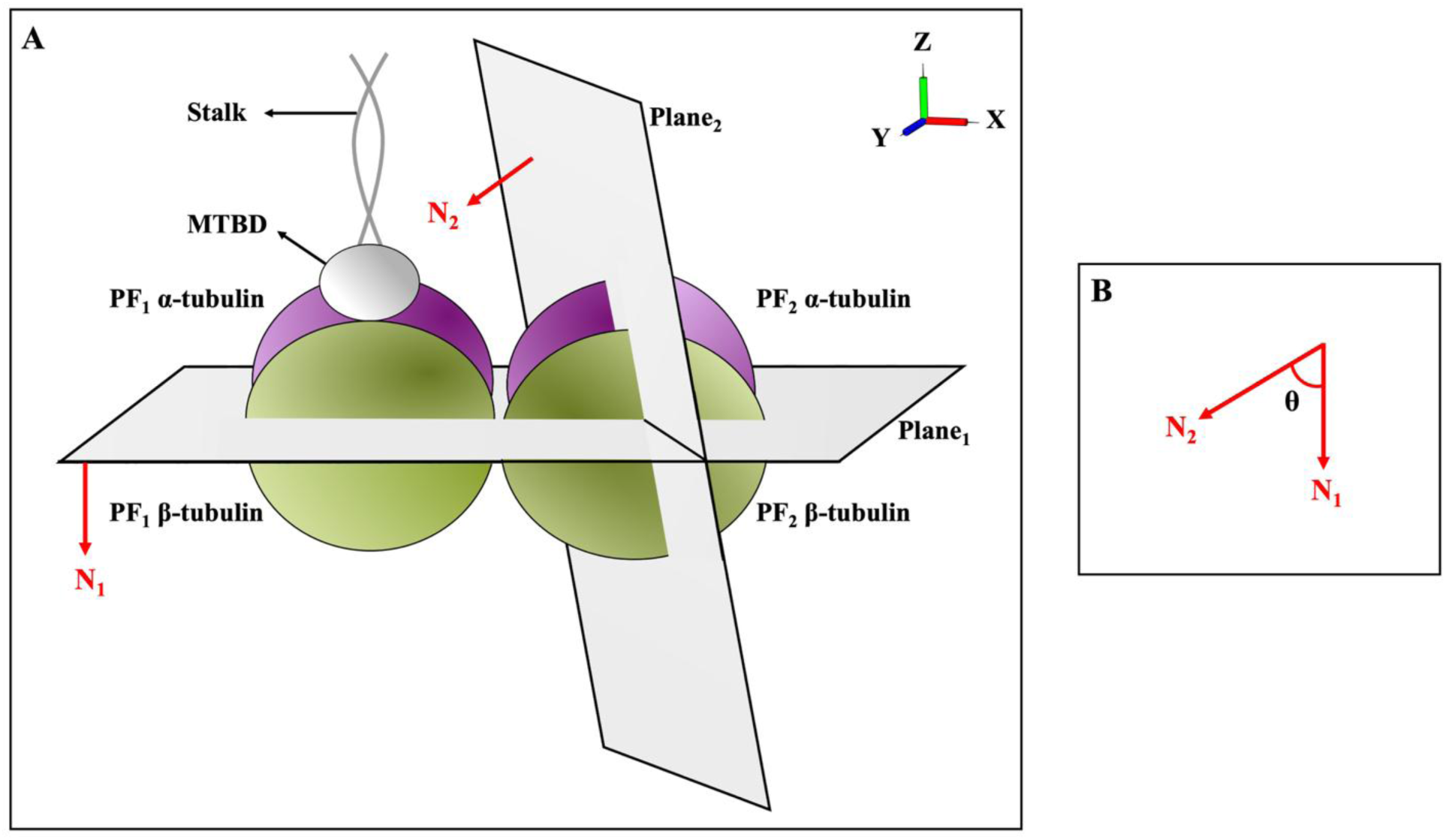
The interdimer angle (θ) at the interface between PF_1_ and PF_2_. (A) The front view of the tubulin protein showing the α- and β-subunits of PF_1_ and PF_2_. Plane _1_ represents the plane through the α, β-tubulin in the PF_1_. Plane_2_ is the plane passing through the α, β-tubulin in the PF_2_.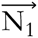 and 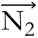 are the normals to Plane_1_ and Plane_2_, respectively, calculated using the singular value decomposition algorithm. (B) Angle θ is the measured angle between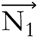 and 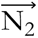 quantifying the deviation of PF_2_ relative to PF_1_.

Once the normal vectors 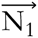 and 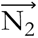 were calculated for a conformation, the interdimer angle (θ) between PF_1_ and PF_2_ was determined by computing the angle between the vectors. (**Eq. 2**)

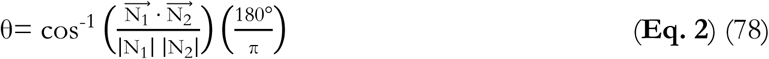

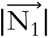 and 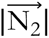 are the magnitudes of the normal vectors and 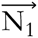. 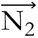 is the dot product of the normal vectors (78).

The interdimer angle θ was calculated for each trajectory frame and plotted against time; it is a quantitative measure of the relative orientation of the PF_2_ with respect to the PF_1_. A θ value less than 90° indicates an anti-clockwise or inward bending of α, β-tubulin of the PF_2_ around the MT’s longitudinal axis towards the α, β-tubulin of PF_1_. In contrast, θ exceeding 90° signifies an outward or clockwise deviation from PF_1_. (**Figure 4**)

## 3. Results and Discussion

The composition, length, and arrangement of amino acids in the CTTs of β-tubulin differ between isotypes. The CTT of wild-type β-tubulin (TUBB3) has a total of 24 amino acids with three (3) aspartates (Asp427, Asp439, Asp440) and nine (9) glutamates (Glu431, Glu432, Glu433, Glu435, Glu438, Glu441, Glu442, Glu443, Glu445), all possessing negative charge. (**Table 2**)

**Table 2:**
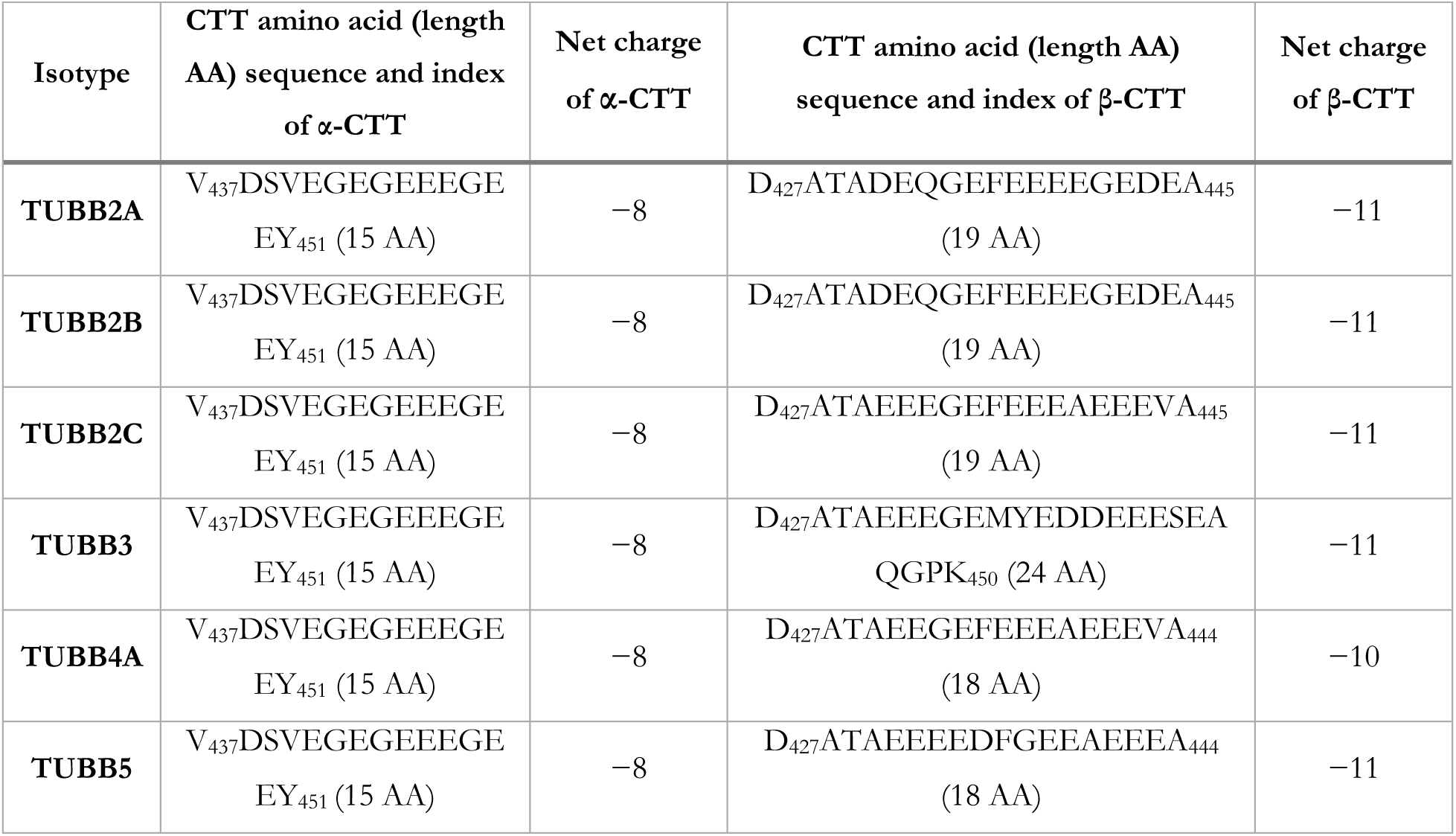
The amino acid (AA) sequence, length, and index of α- and β-CTTs of the tubulin isotypes and total negative and positive charge in each subunit expressed in human brain tumor cell studies in this work.

### 3.1. β-CTT of PF_2_ involves with MTBD bound to TUBB2A, TUBB2B and TUBB2C tetramers

In the tetramer composition, TUBB2A (S7), TUBB2B (S8), and TUBB2C (S9), the β-CTT of the PF_2_ interacts with MTBD bound to the adjacent PF (i.e., PF_1_), lasting from 150 ns to 200 ns of the MD trajectory after convergence. (**Figure S5A–C and Table 1**)

That was not observed in tetramers formed with TUBB3, TUBB4A, and TUBB5 isotypes. (**Figure S5D–F**)

Specifically, in the TUBB2A tetramer (S7), the amino acids of PF_2_’s β-CTT, consisting of Asp443, Glu444, and Ala445, formed hydrogen bonds with the Ser3331 of the MTBD and are involved in several long-range interactions with the MTBD. (**Figure 5A**)

**Figure 5:**
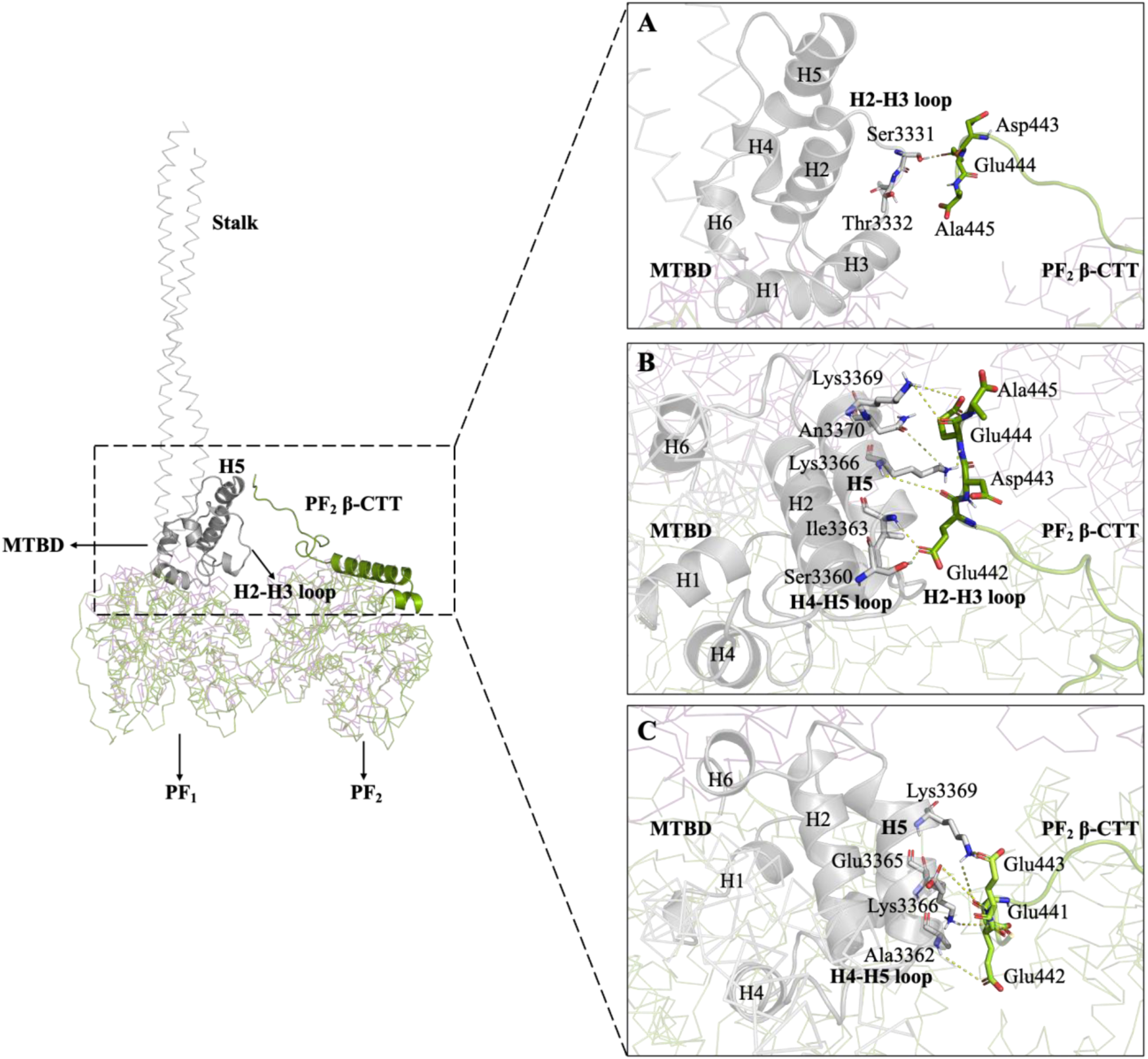
Interactions between the β-CTT of PF_2_ and MTBD in (A) TUBB2A, (B) TUBB2B, and (C) TUBB2C tetramers. β-CTT amino acids are shown as green sticks, and MTBD residues as grey sticks. Key MTBD helices and loops involved in the interactions are labeled in bold.

In the TUBB2B tetramer (S8), Glu440, Glu442, Asp443, and Glu444 of the β-CTT formed salt bridges with Lys3366 and Lys3369 of the MTBD H5, along with various hydrogen bonds between the β-CTT and H2–H3 loop (amino acids 3328–3334), H4–H5 loop (residues 3353–3360), and helix H5 (amino acids 3361–3371) of the MTBD from 150 ns to 350 ns of the trajectory. (**Figure 5B and Table S3**)

Similarly, in the TUBB2C tetramer (S9), β-CTT Glu437, Glu439, Ala440, Glu441, Glu442, Glu443, Val444, and Ala445 formed various salt bridges and hydrogen bonds with the H4–H5 loop (amino acids 3353–3360) and helix H5 (amino acids 3361–3371) of the MTBD from 110–200 ns and 700–800 ns. (**Figure 5C and Table S4**)

While H2–H3 loop forms the MTBD-MT binding interface, structure elements H4–H5 loop and H5 helix make extensive hydrophobic interactions with the distal portion of stalk’s CC2, facilitating the long-range allosteric communication that regulates the dynein activity and registry changes between low and high binding affinity modes (79). Also, helix H5 transmits the conformational changes within the MTBD necessary for dynein to adopt its high-affinity binding state (discussed later in Section 3.5.1). Thus, the β-CTT interacts directly with regions of the MTBD that control its docking geometry and binding strength to the microtubules, potentially affecting the dynein’s motility in an isotype-specific manner.

Additionally, the pattern of β-CTT binding to the MTBD differed by isotype. In TUBB2A, β-CTT formed hydrogen bonds with the MTBD at nearly the middle of the trajectory (∼560 ns). (**Figure S5A**)

In TUBB2B, more frequent interactions (hydrogen bonds and salt bridges) were observed in the first 200 ns of the trajectory (140–340 ns) after convergence. (**Figure S5B**)

In the case of TUBB2C, the interactions were more irregular, with occasional spikes of interactions followed by periods of simulation time without any short or long-range interactions. (**Figure S5C**)

These results indicate that the β-CTT may act as an additional regulatory element in these isotypes, shaping dynein’s motility by modulating its stepping pattern, binding, or processivity (discussed in Section 3.5.1). Conversely, dynein likely encounters a less restrictive track in isotypes lacking this interaction, potentially facilitating faster movement or altered force production. This variability could create functionally distinct MT lattices that can bias motor activity toward specific cargos or subcellular regions, consistent with the tubulin code.

### 3.2. β-CTT of PF_1_ and MTBD-hosting β-tubulins

The β-CTT of PF_1_ in tubulin dimers (S1–S6) as well as in tubulin tetramers (S7–S12) of both wild-type (TUBB3) and isotypes (TUBB2A, TUBB2B, TUBB2C, TUBB4A, and TUBB5) in *Homo sapiens* showed a different interaction pattern compared to other species, such as *Sus scrofa* (26) and *Bos taurus* (24). In *Homo sapiens*, β-CTT of PF_1_ interacted with the β-tubulin core rather than the MTBD bound to the same PF, due to differences in the amino acid sequences in their CTTs. In *Homo sapiens* wild-type and the isotypes, β-CTT of the PF_1_ in the dimers (S1–S6) as well as in tetramers (S7–S12) interacted with several common amino acids, such as Arg213, Lys216, Arg276, Arg282, Lys297, Arg306, His307, Arg309, Lys336, Arg359, and Lys362 of β-tubulin of the PF_1_. Together, these amino acids cluster to form a highly electropositive surface patch on the β-tubulin core, which anchors the negatively charged β-CTT through multiple salt bridges and hydrogen bonds; thereby restricting the mobility of the β-CTT and keeping it tethered to the tubulin surface (i.e., away from MTBD) throughout the trajectory.

The observed interactions of the β-CTT of PF_1_ with Arg213, Lys216, Arg276 (M-loop), Arg 282 (M-loop), Lys297, His307, Arg359, and Lys362 are in addition to those previously reported by Laurin *et al.* for the *Homo sapiens* TUBA1A/TUBB and TUBA1A/TUBB3 isotypes using the Optimized Potential for Liquid Simulations-All atom (OPLS-AA) force field (80). Freedman *et al.* also reported a different kind of interactions in simulations of *Homo sapiens* TUBA4A/TUB isotype using Assisted Model Building with Energy Refinement 03 (AMBER03) force field (81), where the β-CTT engaged with the tubulin core that involved only aromatic and charged amino acids, without any charged-charged amino acid interactions. In contrast, the present study identified several charged-charged interactions between the β-CTT and the β-tubulin core.

The interactions between β-CTT and the tubulin core represent an allosteric mechanism that regulates MT dynamics and, as a result, could influence dynein function. By contacting the tubulin core, β-CTT affects the conformation and stability of the MT lattice that can further affect dynein motility (82, 83). Evidence from yeast mutants lacking the β-CTT has shown that the absence of this region leads to hyperstable astral MTs and excessive dynein-driven sliding, which causes errors in spindle positioning (82). These results suggest that yeast β-CTT acts as an intrinsic destabilizer, preventing excessive stability and maintaining the lattice in a dynamic state (82, 83). A dynein complex engages with this MT track to generate pulling forces for processes such as spindle positioning. As dynein moves, its motor activity further destabilizes the already dynamic MT track, promoting catastrophe and depolymerization. This effectively destroys the track behind the motor, ensuring that each pulling event remains transient and terminates at the correct time and place. This feedback loop prevents excessive or prolonged spindle movements observed in mutants lacking the β-CTT (82). Therefore, β-CTT-core interactions establish the MT instability necessary for negative feedback that terminates dynein-driven force generation, providing a possible mechanism for precise control of mitotic spindle positioning and faithful cell division. Similarly, in *Homo sapiens*, β-CTT regulates MT dynamics in an isotype-specific manner, as substituting TUBB3 β-CTT with the TUBB β-CTT significantly reduced MT assembly rates, the number of assembly events, and the growth length, indicating that the β-CTT sequence encodes regulatory information that dictates MT dynamicity and stability (84). These results suggest that β-CTT-core interactions can alter the conformation and stability of the MT lattice in *Homo sapiens*, which in turn could affect dynein motility.

### 3.3. Isotype-specific lateral interactions modulate the PF geometry in tubulin tetramers

In the studied tubulin isotypes, the amino acid sequence variations lie in two functionally significant regions, either at the lateral interface between the β-tubulins of the adjacent PFs and the intrinsically disordered CTTs. Excluding the CTTs, all the amino acid differences among the studied isotypes cluster around the lateral interface. These amino acid substitutions map across key structural elements, including helices H2, H5–H7, H13, and H14, as well as beta-strands E1, E3–E4, E6–E7, and E9–E11, and their connecting loops. These loops and helices constitute the lateral interface between adjacent β-tubulins, where they participate in hydrogen bonds, salt bridges, and van der Waals interactions that dictate the geometry and strength of inter-protofilament interactions. (**Figure 6 and Table S5**)

**Figure 6:**
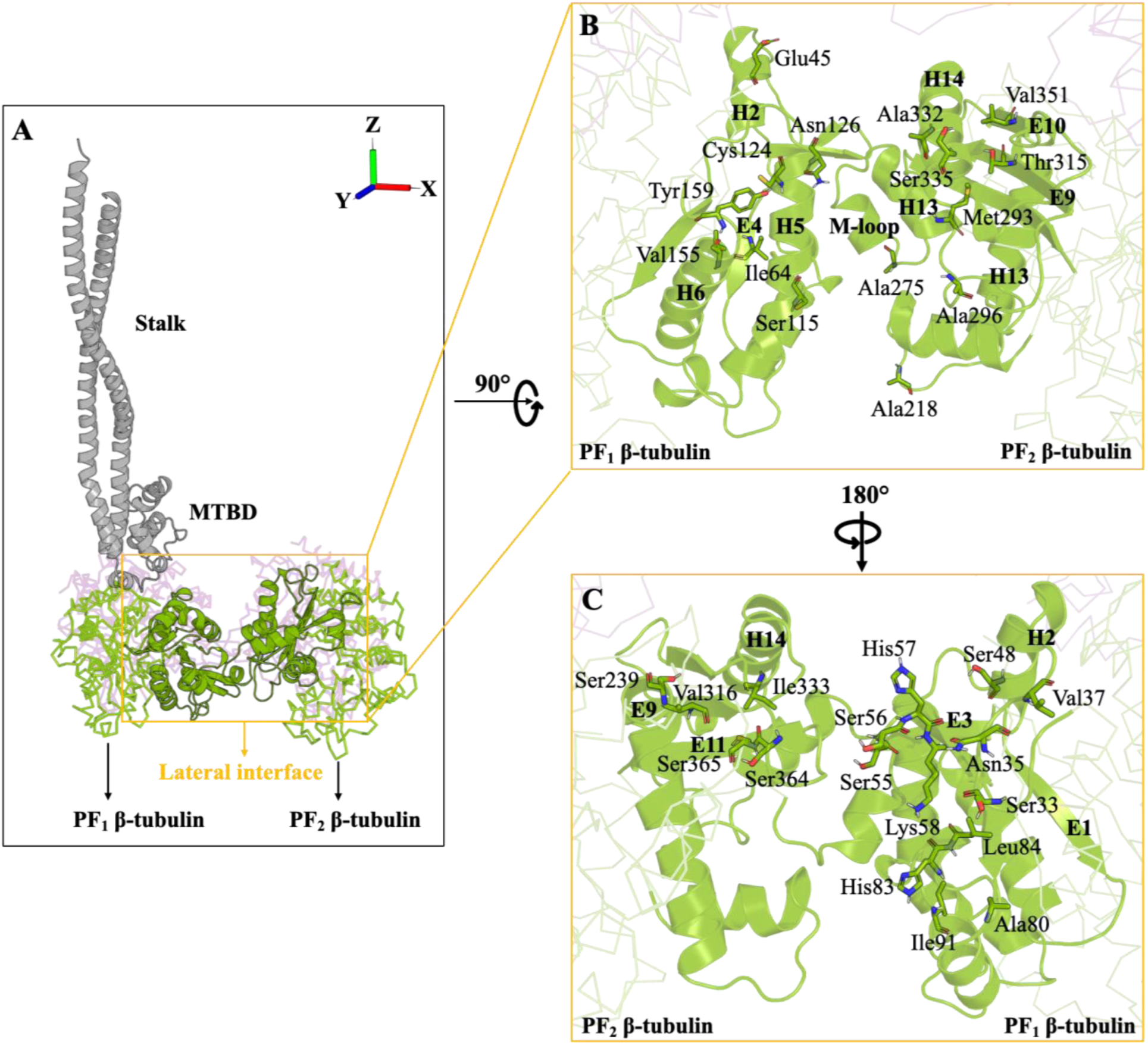
(A) Schematic representation of the tubulin tetramer, showing the lateral interface (in yellow box) where sequence differences between the β-subunits of PF_1_ and PF_2_ are located. (B) and (C) Close-up views of the lateral interface, highlighting the specific amino acids that differ with the corresponding helices and strands indicated in bold. Amino acids are shown for the wild-type tubulin. For corresponding amino acid variations in other isotypes, refer to **Table S5**.

Consistent with these sequence variations at the lateral interface, the isotypes displayed marked differences in the number of long-lasting (occurring for >200 ns after convergence) inter-PF interactions (i.e., between the β-subunits of PF_1_ and PF_2_). Specifically, in each of the TUBB2A, TUBB2B, and TUBB2C isotypes, 1–2 salt bridges and 11–14 hydrogen bonds were identified between adjacent β-subunits, resulting in strong lateral cohesion between PFs (i.e., PF_1_ and PF_2_). In contrast, the TUBB3, TUBB4A, and TUBB5 isotypes lacked long-lasting salt bridges with 8–9 stable hydrogen bonds between the neighboring β-subunits. (**Figure 7 and Table S6**)

**Figure 7:**
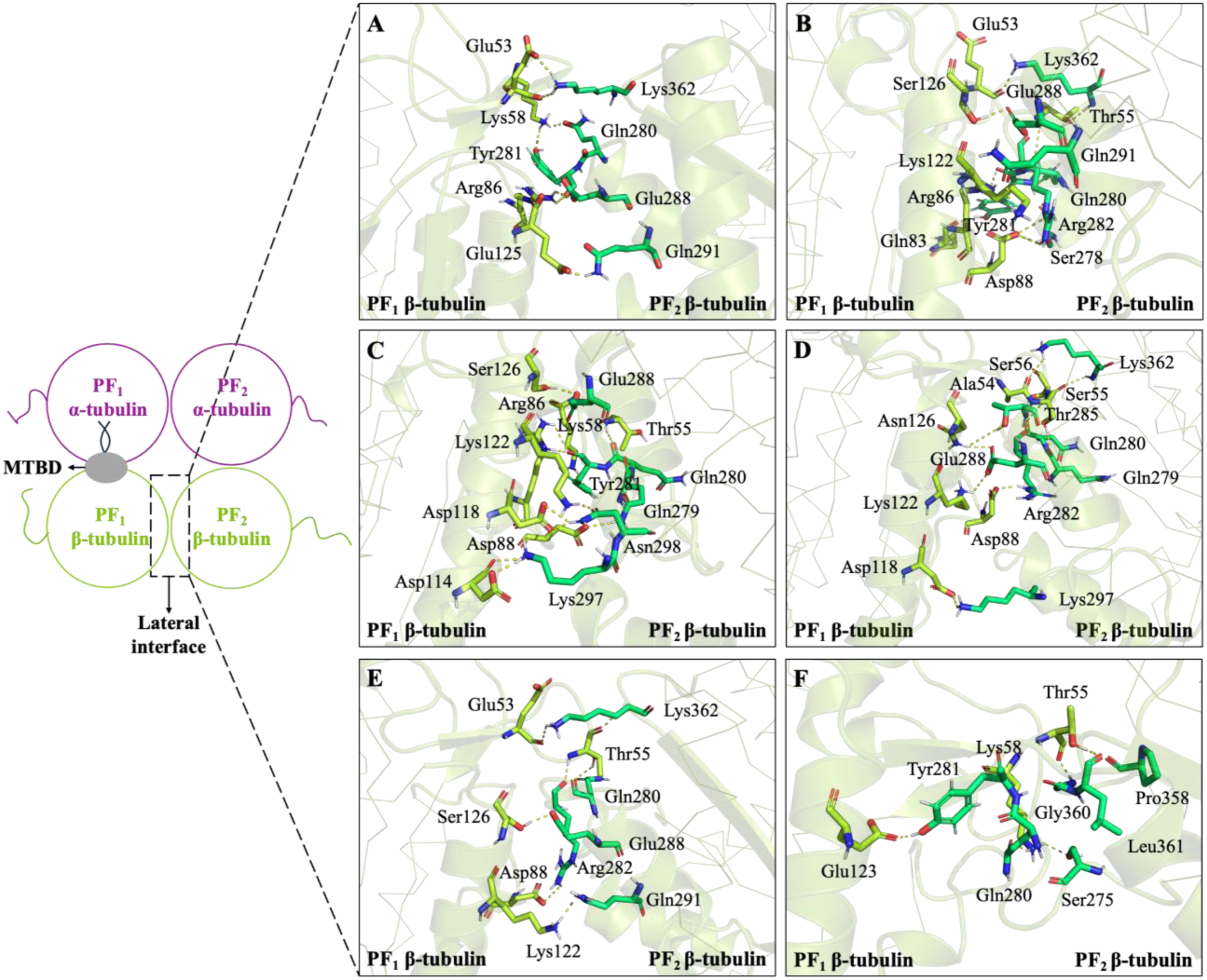
Interacting amino acids at the lateral interface between the β-tubulin subunits of PF_1_ (light green sticks) and PF_2_ (teal sticks) in (A) TUBB2A, (B) TUBB2B, (C) TUBB2C, (D) TUBB3, (E) TUBB4A, and (F) TUBB5 isotypes. The M-loop consists of 13 amino acids from Pro272 to Leu284. A complete list of interacting amino acids is provided in **Table S6**.

This variation in the number of interactions between the adjacent β-subunits was also reflected in the non-bonded interaction energy (electrostatics and van der Waals) between the PF_1_ and PF_2_. TUBB2A (−1012.04 kJ/mol), TUBB2B (−1316.26 kJ/mol), and TUBB2C (−1252.68 kJ/mol) exhibited significantly lower (i.e., more favorable) interaction energies than TUBB4A (−944.37 kJ/mol) and TUBB5 (−1000.25 kJ/mol), indicating stronger inter-PF interactions in the former. Interestingly, TUBB3, despite lacking long-lasting salt bridges, exhibited a relatively strong interaction energy of −1307.84 kJ/mol, comparable to that of TUBB2B and TUBB2C. This can be attributed to two charged histidine residues (His57 and His83) in TUBB3, which are uncharged amino acids (Asn, Gln, or Gly) in the other isotypes. These histidines are involved in multiple short-lived (<10% of the trajectory time) electrostatic interactions at the lateral interface, thereby contributing to the overall stabilization between PF_1_ and PF_2_ in TUBB3.

Crucially, these differences in the lateral interactions among isotypes are directly translated into PF geometry, as indicated by the changes in the interdimer angle between laterally associated PFs. Isotypes with stronger lateral interactions, TUBB2A (1 salt bridge and 11 hydrogen bonds), TUBB2B (2 salt bridges and 14 hydrogen bonds), and TUBB2C (1 salt bridge and 14 hydrogen bonds), exhibited relatively constrained interdimer angle fluctuations, ranging from 20° to 80°. In contrast, isotypes with weaker lateral interactions, such as TUBB3 (9 hydrogen bonds), TUBB4A (9 hydrogen bonds), and TUBB5 (8 hydrogen bonds), exhibited broader fluctuations in the lateral interdimer angle, ranging from 30° to 120°, indicating reduced lateral stability and greater PF flexibility. (**Figure S6**)

A possible explanation for these variations in the interdimer angle is the intrinsic ability of tubulin isotypes to assemble into MTs with different protofilament numbers. Previous studies have shown that human TUBB2B and TUBB3 isotypes can form MTs with distinct PF numbers (21), and such variations in PF number introduce geometric mismatches in the lattice (85). To accommodate these mismatches, lateral loops and helices undergo subtle conformational adjustments (21), which in turn alter the interdimer angle between the PFs. This agrees with recent cryoelectron microscopic data, which showed that MTs with different PF numbers have different inter-protofilament angles (86).

Together, these results demonstrate how tubulin isotypes fine-tune MT lattice biomechanics. Variations in lateral interactions among isotypes alter PF bending rigidity, which in turn governs lattice plasticity and stability. Experimental data support this interpretation; yeast tubulin, enriched in polar residues at the lateral interface, assembles into MTs with altered protofilament numbers compared to those of mammalian tubulin (87). Likewise, Ti *et al.* showed that *Homo sapiens* αIB/βIIB-tubulin supports a broader distribution of PF numbers and greater lattice plasticity than αIB/βIII, which preferentially stabilizes 13-PF microtubules (21). Our findings align with these observations, suggesting that specific isotypes act as lattice modulators, tuning the balance between rigidity and flexibility in the MT lattice. Such modulation has profound implications for dynein’s function. Dynein’s motility is sensitive to MT geometry and deformation. Previous experiments have shown that dynein transport is facilitated by buckled or strained MTs in a low-strain (≤20%) elastic regime, where lattice deformation primarily affects inter-PF interactions (88). Thus, isotypes such as TUBB3, TUBB4A, and TUBB5, with weaker lateral stabilization and greater PF flexibility, may form MT tracks that are more easily deformed and therefore more permissive to dynein motility. In contrast, isotypes like TUBB2A, TUBB2B, and TUBB2C, with strong lateral cohesion, may support more stable tracks that could facilitate spindle movement by promoting dynein-driven lateral sliding either by providing more accessible binding sites for dynein to traverse along their lateral sides or allowing them to withstand better cortical stresses such as bending and the forces generated by dynein motility (82). This duality suggests that cells use isotype-specific lateral interactions and PF assembly properties to regulate the cytoskeleton’s architecture and functional specialization.

### 3.4. The interprotofilament angle (θ) affects the β-CTT-MTBD interactions

The calculated lateral interdimer angles between the adjacent PFs exhibited a direct relationship with the distance between the center of mass (dCOM) of the MTBD bound to the PF_1_ and the β-CTT in PF_2_ in tubulin tetramers. Smaller interprotofilament (i.e., interdimer) angles correlated with closer β-CTT-MTBD proximity, whereas larger angles correlated with separation, indicating that lateral bending of PFs modulates how closely the β-CTT can approach the MTBD on the adjacent PF.

Isotypes (TUBB2A, TUBB2B, and TUBB2C) in which the β-CTT of PF_2_ interacts with the MTBD showed consistently smaller angles (TUBB2A: 35°–85°, TUBB2B: 25°–65°, TUBB2C: 25°–75°) and correspondingly short dCOM distances (TUBB2A: 30–75Å, TUBB2B: 20–60Å, TUBB2C: 20–70Å). In contrast, TUBB3 (55°–95°), TUBB4A (30°–90°), and TUBB5 (60°–120°) sampled larger interdimer angles and exhibited increased dCOMs (TUBB3: 50–80Å, TUBB4A: 50–90Å, TUBB5: 55–90Å) between the β-CTT of PF_2_ and the MTBD. (**Figure 8**)

**Figure 8:**
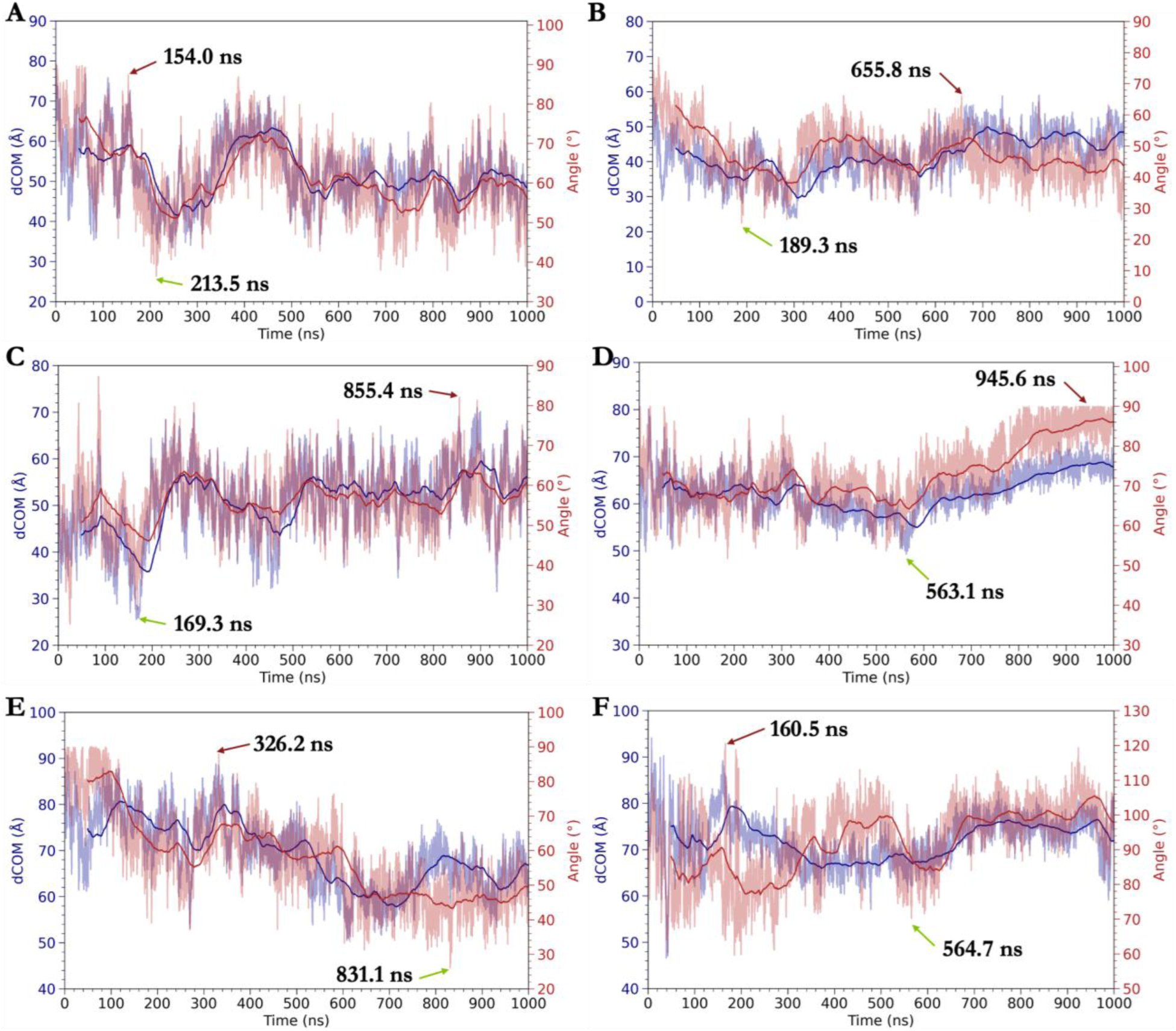
Correlation between dCOM and the interdimer angle for (A) TUBB2A (S7), (B) TUBB2B (S8), (C) TUBB2C (S9), (D) TUBB3 (S10), (E) TUBB4A (S11), and (F) TUBB5 (S12) tetramers. The dCOM was calculated between the MTBD bound to the PF_1_ and the β-CTT of the PF_2_. The interdimer angle is determined between the two neighboring PFs (PF_1_ and PF_2_). In each plot, the blue line represents the progression of dCOM, while the red line illustrates the variation in the interdimer angle over time. The solid lines depict the 500-point moving average, highlighting the overall trend and correlation between the dCOM and lateral interdimer angle throughout the trajectory. The green and red arrows represent the conformations with the lowest and highest interdimer angle, respectively.

This increase in the β-CTT and MTBD distance arose from a sequence of conformational changes in the β-tubulin of the PF_2_. Increased interdimer angle (ranging from 30°–120°) in the TUBB3, TUBB4A, and TUBB5 isotypes facilitated the outward rotation (away from the PF_1_) of β-tubulin of the PF_2_ around the MT longitudinal axis, consistent with experimental evidence that non-13-PF microtubules accommodate altered lattice geometry through either PF sliding (i.e., lattice shear) or rotation around the MT’s long axis (i.e., lattice rotation) (85). (**Figure 9**)

**Figure 9:**
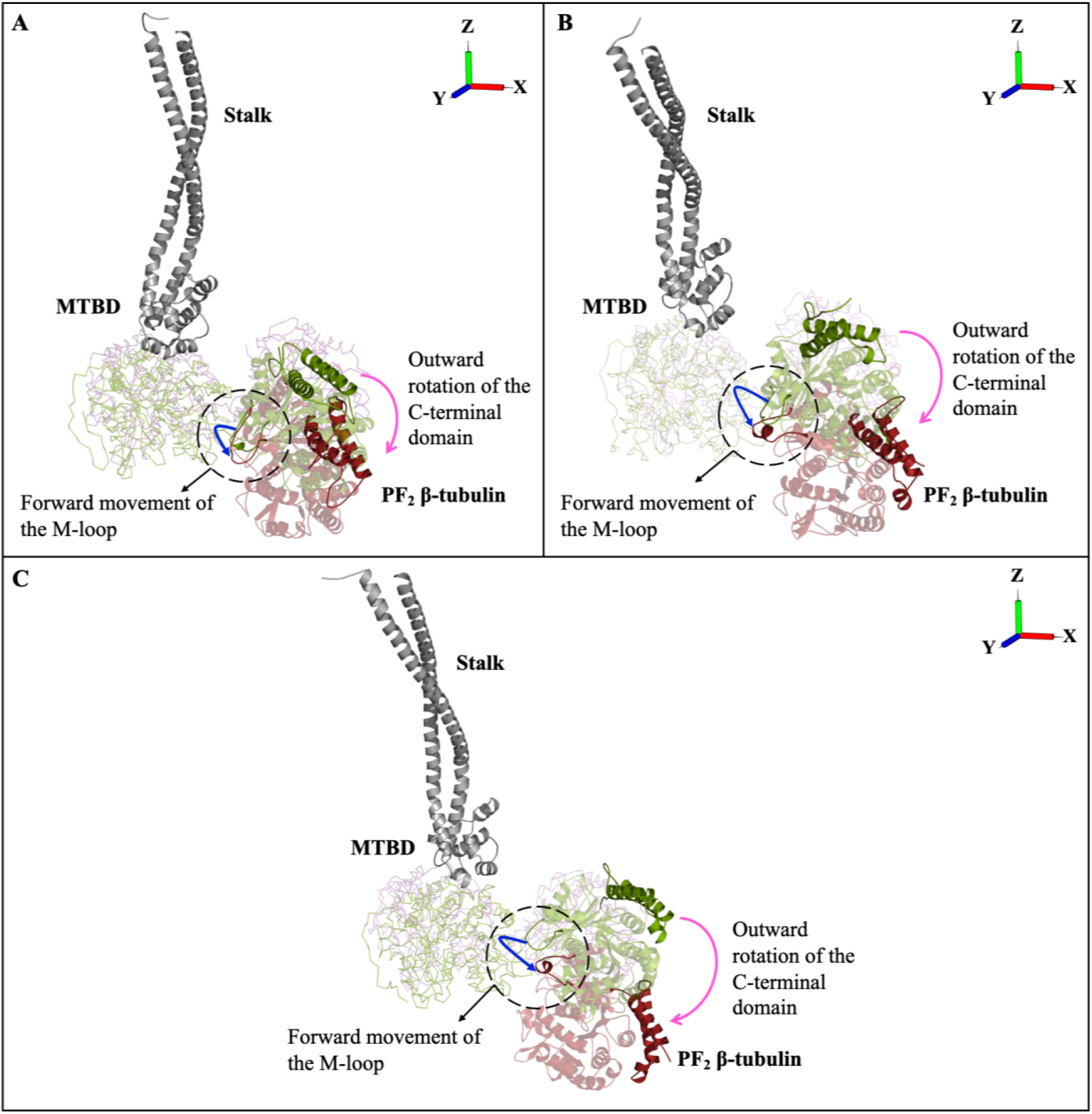
Schematic showing the rotation of β-tubulin in PF_2_ around the MT’s longitudinal axis due to interdimer bending relative to PF_1_, accompanied by the forward movement of the M-loop (blue arrows) and the outward (i.e., away from the PF_1_; pink arrows) displacement of C-terminal helices and the β-CTT of the PF_2_ (shown in cartoon) away from the MTBD in the (A) TUBB3, (B) TUBB4A, and (C) TUBB5 tetramers. Green and red conformations correspond to the lowest and highest interdimer angles for each isotype, as indicated in Figure 8.

This outward rotation (i.e., away from the PF_1_) of the PF_2_ in the TUBB3, TUBB4A, and TUBB5 isotypes caused a conformational shift in the M-loop of the PF_2_’s β-tubulin. Initially interacting with Asp88 of β-tubulin of PF_1_, the M-loop shifted forward to establish new interactions with Glu123, Glu125, and Asp128 of helix H5 of β-tubulin in the PF_1_ in these isotypes. (**Figure 9**)

These interactions were absent in TUBB2A, TUBB2B, and TUBB2C, where the strong lateral interactions (1–2 salt bridges and 11–14 hydrogen bonds) prevented the conformational changes required for this shift. Hydrogen bond analysis supports these findings: in TUBB3, a new hydrogen bond between Thr285 (M-loop of PF_2_) and Asn126 (β-subunit of PF_1_) forms over 35% of the trajectory time (i.e., occupancy). (**Figure 7D**)

In TUBB4A, where the interdimer angles were intermediate (ranging from 30° to 90°), a similar interaction between Ser126 of PF_1_ and Thr285 of the PF_2_ M-loop was observed, but with much lower occupancy (i.e., <5% of the trajectory time). In TUBB5, which had the widest interdimer angles (ranging from 60° to 120°), a strong interaction was observed between Tyr281 (M-loop of PF_2_) and Glu123 (H5 of PF_1_), with over 80% of the trajectory time (i.e., occupancy), highlighting the structural rearrangement enabled by the more flexible lateral interface. (**Figure 7F**)

Such M-loop rearrangements propagated structural changes to C-terminal helices (H15, H16, and H17), which shifted outward (i.e., away from the PF_1_), displacing the β-CTT away from the MTBD in the TUBB3, TUBB4A, and TUBB5 isotypes. As a result, the β-CTT of PF_2_ could not interact with the MTBD, in contrast to TUBB2A, TUBB2B, and TUBB2C, where a tight lateral packing due to the more lateral interactions (№. 1–2 salt bridges and №. 11–14 hydrogen bonds) preserved proximity between the β-CTT of PF_2_ and the MTBD of PF_1_. (**Figure 9**)

These findings establish a direct structural pathway through which isotype-specific lateral interactions impact the accessibility of β-CTT of the adjacent PF to the dynein’s MTBD. The β-CTT is recognized as a regulatory element of MT-motor communication, serving as an electrostatic interaction hotspots that influence dynein binding affinity and processivity (30). By modulating interdimer angles, isotypes act as gatekeepers of β-CTT positioning; strong lateral interactions (TUBB2A, TUBB2B, and TUBB2C) maintain β-CTT-MTBD proximity, whereas weak lateral interfaces (TUBB3, TUBB4A, and TUBB5) displace β-CTT away from the MTBD.

This mechanism explains apparently two distinct experimental findings: (i) dynein requires CTTs for optimal binding as CTT truncation strongly reduces dynein motility (30, 34), and (ii) dynein motility can also be facilitated by mechanically deformed lattices (88). The current study provides a mechanistic explanation. When isotypes have small interdimer angles (20°–80°) and maintain β-CTT proximity, dynein benefits from direct CTT-MTBD electrostatic engagement (promoting a high-affinity state); however, when isotypes allow large angles (30°–120°), β-CTTs are displaced, but the lattice becomes more deformable, which, under some conditions (e.g., under force or buckling), may assist dynein stepping and motility even in the absence of direct CTT interactions.

This dual regulation is likely to have tissue-specific consequences. In neurons, where TUBB3/TUBB4A is abundant, lattice flexibility may facilitate long-range dynein-driven cargo transport along highly dynamic MTs. In contrast, in dividing cells, isotypes with stronger lateral interfaces in TUBB2A, TUBB2B, TUBB2C (due to higher inter-PF interactions; 1–2 salt bridges and 11–14 hydrogen bonds) as compared to TUBB3, TUBB4A, and TUBB5 may produce stable and stiffer MTs that have been previously reported to faclitate spindle movement by promoting dynein-driven lateral sliding (82). Indeed, TUBB2C is among the most highly expressed β-tubulin isotypes in dividing cells (16), comprising up to 22% of total β-tubulin (89). Similarly, mutations in TUBB2A (89) or TUBB2B (90) impair spindle pole assembly and delay mitotic progression. Together, these studies highlight the essential role of these isotypes (TUBB2A, TUBB2B, and TUBB2C) in cell division, possibly due to their higher inter-PF interactions (№. 1–2 salt bridges and №. 11–14 hydrogen bonds), leading to stabilization of MTs. Changes in isotype composition in disease (e.g., TUBB3 upregulation in cancer) could therefore change intracellular transport, mitotic stability, and anticancer drug response by altering the lattice geometry and β-CTT accessibility.

### 3.5. β-CTT of the PF_2_, conformational changes in MTBD, and their large-scale collective motions

Since in TUBB2A, TUBB2B, and TUBB2C isotypes, the β-CTT of PF_2_ interacts with the MTBD and also, these isotypes are known to be involved in cell division as discussed in the above section (Section 3.4), these isotypes were further subjected to Principal Component Analysis (PCA) to characterize their global collective motions and elucidate the impact of β-CTT of PF_2_ on the conformational changes in the MTBD. PCA showed that the essential motions of tubulin tetramers differ among the TUBB2A, TUBB2B, and TUBB2C isotypes. In TUBB2A, the first three principal components (PC) accounted for 77.56% of the total variance, with contributions of 45.34% (PC1), 26.71% (PC2), and 5.51% (PC3). For TUBB2B, PC1, PC2, and PC3 contributed 64.1%, 22.37%, and 3.15% of the variance, respectively, representing a cumulative 89.62%. In TUBB2C, the first three principal components together explained 65.42% of the variance, with contributions of 35.43% (PC1), 23.24% (PC2), and 6.75% (PC3). These values indicate that the essential subspace of the system dynamics is well represented by the top three PCs, justifying their selection for further analysis. (**Figure 10A–C**)

**Figure 10:**
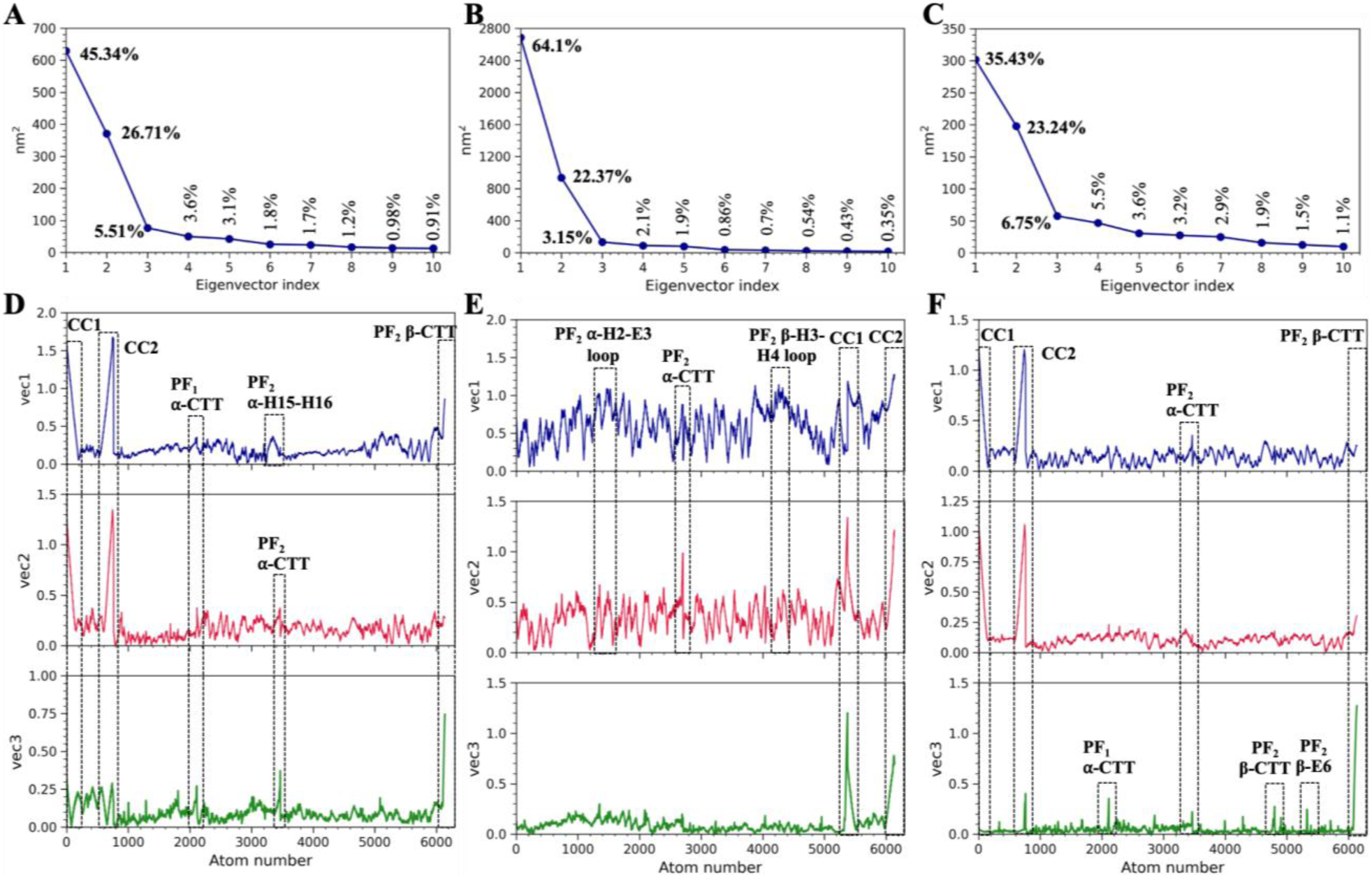
Scree plot of eigenvalues (nm^2^) vs eigenvector index for the first 10 modes with contribution of each mode of (A) TUBB2A, (B) TUBB2B, and (C) TUBB2C isotypes. The root mean square fluctuations (RMSF) of the backbone atoms for the first three eigenvectors of (D) TUBB2A, (E) TUBB2B, and (F) TUBB2C isotypes.

The cosine content of the first three principal components (PC1–PC3) was ≤0.4 across all isotypes, confirming that the extracted motions are not dominated by random Brownian diffusion. A high cosine content (>0.7) reflects stochastic noise or insufficient sampling, while lower values (≤0.5) signify well-converged sampling and genuine collective motions (91). Thus, the collective motions identified in this study represent meaningful conformational transitions rather than artifacts of diffusion. The trace of the covariance matrix was highest for TUBB2B (4191.41 nm²), compared to 1389.28 nm² for TUBB2A and 851.85 nm² for TUBB2C, indicating that TUBB2B explores a more extensive conformational space. Consistent with this, projection of the MD trajectories onto the first three principal components (PC1–PC2, PC2–PC3, and PC1–PC3) further highlighted isotype-specific differences. TUBB2B exhibited a broader distribution and distinct clustering in phase space, whereas TUBB2A and TUBB2C sampled narrower conformational ranges. (**Figure S7**)

The dominant collective motion captured by the first and second eigenvectors in TUBB2A, TUBB2B, and TUBB2C isotypes involves a large rocking and twisting motion of the stalk domain, which is transmitted into displacements of the MTBD at its binding site on PF_1_. (**Figure 11**)

**Figure 11:**
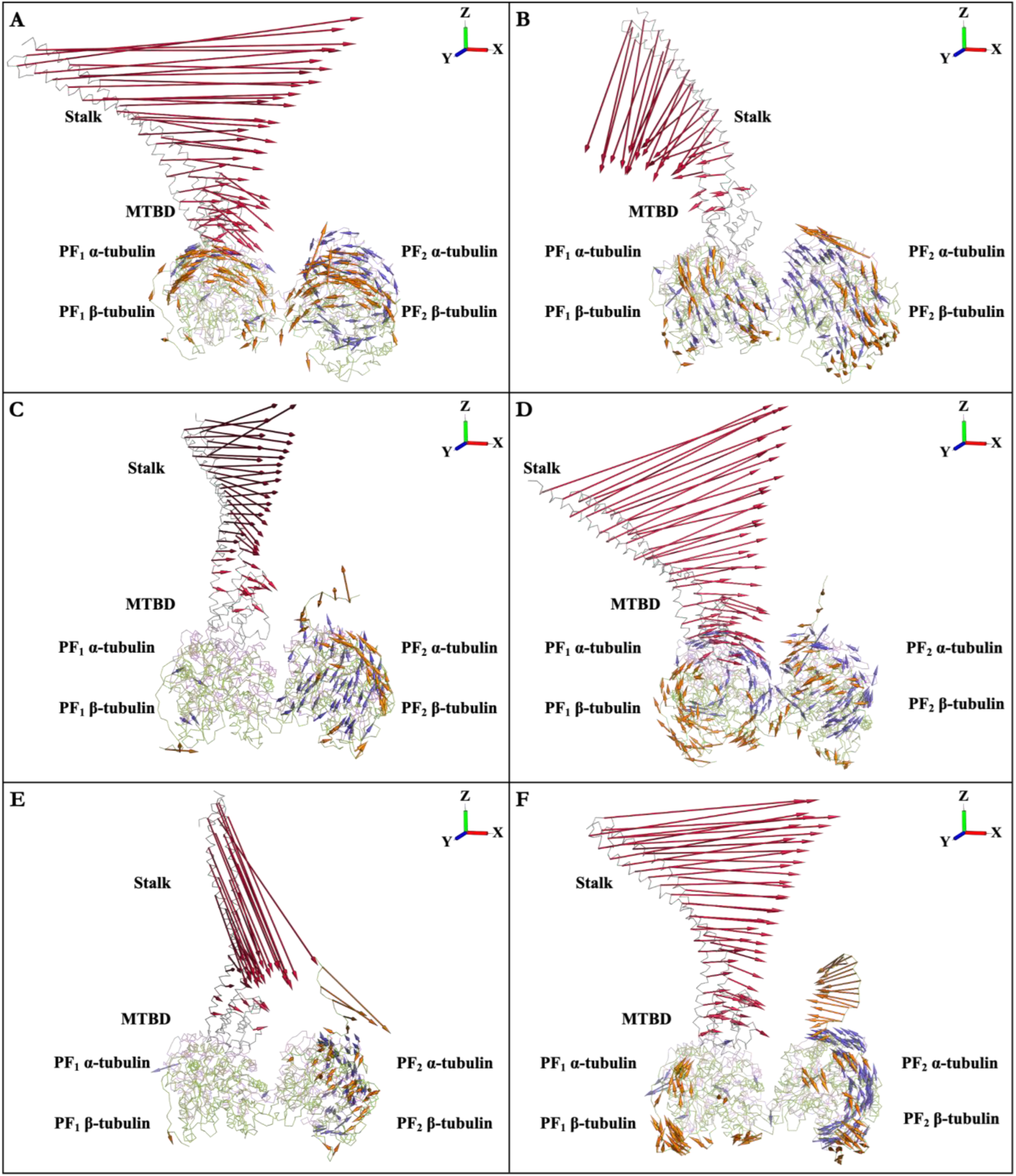
Plots depicting the dominant motions along the first eigenvector for (A) TUBB2A, (C) TUBB2B, and (E) TUBB2C, and along the second eigenvector for (B) TUBB2A, (D) TUBB2B, and (F) TUBB2C isotypes. Length of arrows indicates magnitude of the motion: red for MTBD and stalk, blue for α-tubulins of PF_1_ and PF_2_, and orange for β-tubulins of PF_1_ and PF_2_.

Importantly, this motion is coupled to the movement of the β-CTT of PF_2_ toward or away from the MTBD, consistent with the eigenvector-projected root-mean square fluctuations (RMSF), which show the largest amplitudes in the stalk coiled coils and the β-CTT of PF_2_ in the PC1 and PC2 in these isotypes. (**Figure 10D–F**)

Additionally, the α- and β-subunits of the PF_2_ showed movement towards the PF_1_, consistent with the local interactions in the preceding sections, indicating a stronger lateral cohesion and reduced interdimer angle between the adjacent PFs in these isotypes. (**Figure 11**)

The Dynamic Cross-Correlation (DCC) analysis provided further insight into how the dynein stalk/MTBD system is mechanically coupled to PF_2_’s β-CTT in TUBB2A, TUBB2B, and TUBB2C isotypes. The maps showed strong anti-correlated motions between α- and β-tubulin of the PF_2_ and the MTBD, as well as with the stalk domains CC1 and CC2 in these isotypes. Anti-correlation in this context means that the two regions move in opposite directions but in a coordinated way. When the MTBD/stalk shifts toward PF_2_, the β-CTT of PF_2_ is simultaneously displaced toward the MTBD, bringing the two surfaces closer together. This synchronized but opposing displacement effectively reduces the spatial gap between the PF_2_ β-CTT and the MTBD on PF_1_, enhancing the likelihood of transient contacts between them. (**Figure 12**, **Figure 11A, C–D, and F**)

**Figure 12:**
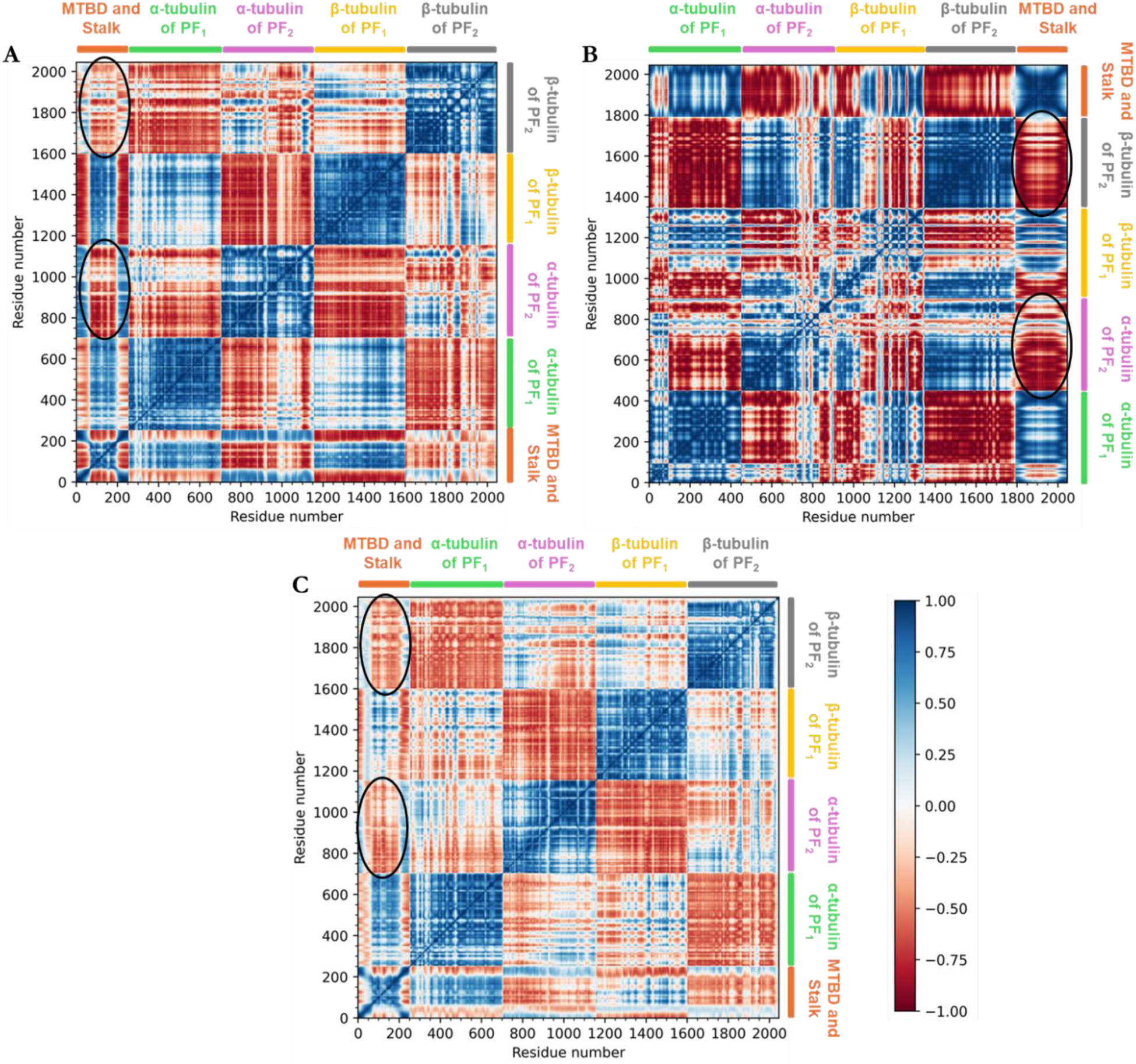
Dynamic Cross-Correlation maps of (A) TUBB2A, (B) TUBB2B, and (C) TUBB2C isotypes. The extent of correlated and anti-correlated motions is color-coded, blue to white to red. Blue indicates the motions in the same direction, while red indicates motions in the opposite direction. The marked black circles represent the anti-correlated motions of α, β-tubulin heterodimer in the PF_2_ and the MTBD and stalk domain.

This agrees with the PCA, where the dominant mode involved the β-CTT approaching the MTBD, and with the eigenvector-projected RMSF, which revealed that both the stalk and PF_2_ β-CTT contribute significantly to dominant motions. (**Figure 11A, C–D, and F and Figure 10D–F**)

Conversely, when the MTBD/stalk moves away from PF_2_, the β-CTT also withdraws, increasing their separation. Together, these results indicate that β-CTTs are dynamically steered toward or away from the MTBD in coordination with stalk-driven rocking, providing a structural mechanism for regulating dynein-microtubule engagement. (**Figure 11**)

These anti-correlated motions between the dynein (i.e., MTBD and stalk) and the α, β-tubulin heterodimer in PF_2_ indicate that the lateral interface (i.e., M-loop) between the PFs (i.e., PF_1_ and PF_2_) acts as a hinge, in agreement with the recent cryoelectron microscopy study of bovine MTs that have shown that adjacent PFs pivot around their lateral interface, functioning like a hinge with the axis of rotation located primarily at the M-loop (86). This intrinsic flexibility enables PFs to move toward or away from each other, which brings the MTBD and the β-CTT of PF_2_ into alternating proximity and separation, consistent with the observed DCC patterns. (**Figure 12**)

Electrostatic complementarity between the MTBD/stalk and β-tubulin of PF_2_ further contributes to these anti-correlated motions in TUBB2A, TUBB2B, and TUBB2C isotypes. The β-H17 region and β-CTT of PF_2_ are enriched with acidic amino acids (Glu405, Glu407, Glu410, Glu412, Asp417, Glu421, Asp427, Glu432, Glu435, Glu437, Glu438, Glu439, and Glu442), creating a highly negative electrostatic surface that extends outward from the PF_2_. Aromatic residues (Phe408, Tyr422, Tyr425, and Phe436) further reinforce this negative field, contributing additional electron density due to their partial electron-rich character. (**Figure S8A–B**)

In contrast, the MTBD helices (H2, H3, and H5) and the adjacent stalk domains (CC1 and CC2) that face PF_2_ are populated with lysine, arginine, and histidine residues (Lys3318 in helix H2, Arg3364, Lys3336, Arg3339, and Arg3344 in helix H3, Lys3366, Lys3368, and Lys3369 in helix H5, as well as His3265, Lys3266, Lys3274, Lys3279, and Lys3284 in the CC1, and Lys3407, Arg3408, Arg3413, Lys3418, and Lys3424 in CC2), forming a strongly electropositive surface. This opposing charge distribution creates a long-range electrostatic attraction between the β-CTT of PF_2_ and the MTBD/stalk domain. (**Figure S8C–D**)

Consequently, when the β-CTT is in an unbound conformation (i.e., not engaged with the β-tubulin surface of PF_2_), it is drawn toward the positively charged MTBD and stalk domain, reflecting an electrostatic steering mechanism that dynamically positions the β-CTT of the PF_2_ toward the MTBD, favoring transient interactions between them. This conclusion aligns with a previous study indicating that β-CTT guides the MTBD to its binding position on the α, β-tubulin heterodimer via long-range electrostatic interactions (24). (**Figure 11A, C–D and F**)

Further, this interpretation is consistent with the PCA results, where the first two eigenvectors captured the movement of the β-CTT of the PF_2_ towards the MTBD, and with the eigenvector-projected RMSF, which highlighted significant contributions from both the stalk coiled coils and PF_2_’s β-CTT. (**Figure 11A, C–D and F and Figure 10D–F**)

#### 3.5.1. β-CTT-mediated modulation of the MTBD conformation across tubulin isotypes

The highest variance motion captured by the first eigenvector revealed an isotype-specific mechanism by which the β-CTT of an adjacent PF regulates motor binding affinity. In the TUBB2A, TUBB2B, and TUBB2C isotypes, β-tubulin of PF_2_ moved closer to PF_1,_ which brought the β-CTT into the vicinity of the MTBD. In this position, the β-CTT interacted with the H2–H3 loop, the H4–H5 loop, and helix H5 of the MTBD, resulting in local structural adjustments consistent with the local interactions. (**Figure 5**)

In TUBB2A, the conformations extracted along the first eigenvector showed that MTBD helix H5 rotated by 40° (in conformation 1) and 10° (in conformation 2) in the YZ plane, accompanied by translational shifts of 17.6 Å (in conformation 1) and 2.53 Å (in conformation 2) as compared to the initial frame of the trajectory due to its interactions with the β-CTT of PF_2_. These structural adjustments in the MTBD H5 were transmitted into the neighboring H5–H6 loop, leading to the reorientation of MTBD helix H6 and shifting it closer to the α-tubulin of PF_1_, enabling Ser3376 (H5–H6 loop), Asn3383, and Arg3384 of MTBD helix H6 to interact with α-tubulin H14–H15 loop (Tyr399, Arg402) and helix H16 (Glu415) for up to 30% of the simulation time (i.e., occupancy). (**Table S7 and Figure S9**)

In TUBB2B, the MTBD H5 rotated by 29.5° (in conformation 1) and 15.6° (in conformation 2) on the YZ surface and shifted by 7.6Å (in conformation 1) and 9Å (in conformation 2) as compared to the initial frame of the trajectory. These displacements are driven by interactions with the β-CTT of PF_2_ and propagated into the neighboring H5–H6 loop, leading to a shift of MTBD helix H6 toward the α-tubulin of PF_1_. This repositioning of H6 enabled Ser3376 (H5–H6 loop), Asn3383, and Arg3384 of MTBD H6 to interact with H14–H15 loop (Tyr399, Arg402) and helix H16 (Glu415, Ser419) of the α-tubulin of PF_1_ for 10–55% of the simulation time (i.e., occupancy). (**Table S7 and Figure 13**)

**Figure 13:**
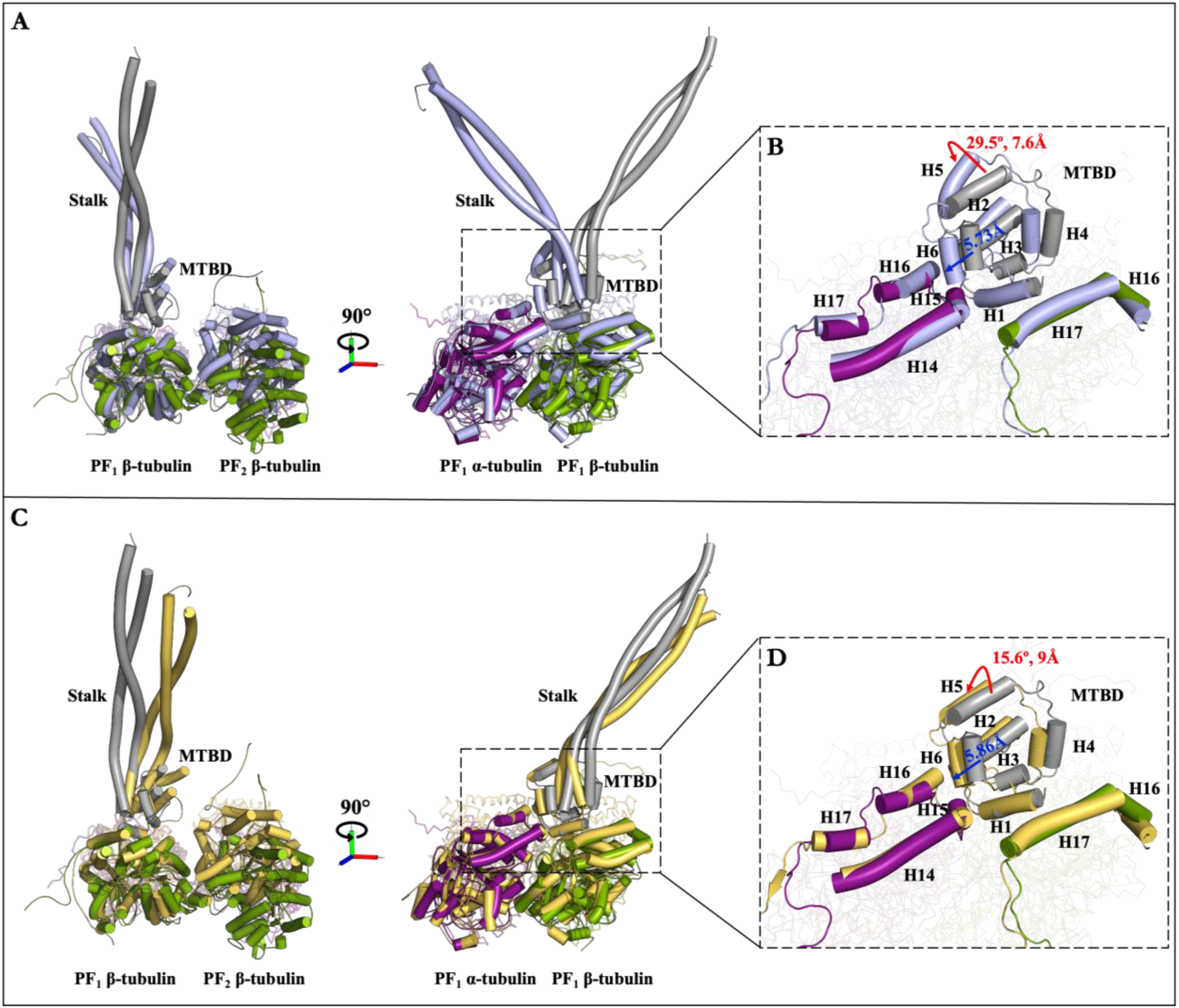
Comparison of two distinct conformations of the TUBB2B tetramer from the first principal component analysis (PCA) (light blue and orange yellow) with the initial trajectory frame (α-tubulin: purple, β-tubulin: green, MTBD/stalk: gray). Panels (A) and (C) show the overall conformational differences of conformation 1 (light blue) and conformation 2 (orange yellow), respectively, while panels (B) and (D) present close-up views of the MT-MTBD interface in these two conformations with red and blue arrows showing the movement of MTBD helices H5 and H6, respectively.

In TUBB2C, the MTBD H5 rotated by 22.7° and 12.3° and shifted by 3.63Å and 6.2Å in conformation 1 and 2, respectively, compared to the first frame of the trajectory, due to its interactions with the β-CTT of PF_2_. These changes in the MTBD H5 were propagated to helix H6 through the H5–H6 loop. Helix H6 moved closer to α-tubulin of PF_1_ by 7.75Å (in conformation 1) and 6.9Å (in conformation 2), where Arg3384 and Arg3385 of MTBD H6 started interacting with the helix H16 (Glu414, Glu415) of the α-tubulin of PF_1_ for 20–52% of the simulation time (i.e., occupancy). (**Table S7 and Figure S10**)

MTBD H6, together with helices H1 and H3, is a key element of the high-affinity binding mode of the MTBD. Structural and mutagenesis studies in yeast (*Saccharomyces cerevisiae*) and mice dynein have indicated that the transition from the low to high binding mode requires MTBD H6 to move closer to the α-tubulin, and the interactions between the MTBD H1, H3, and H6 and MT are required to induce and maintain this high-affinity conformation (11, 32, 92). Likewise, mutating α-tubulin residues (Arg403 and Glu416 in *Saccharomyces cerevisiae*; corresponding to Arg402 and Glu415 in *Homo sapiens*) that interact with the MTBD H6 abolishes dynein motility and are critical for dynein to switch from a weak to a strong binding state (45). While helix H5 does not directly interact with the MT, it transmits the conformational changes induced by β-CTT of PF_2_ through the H5–H6 loop into a reorientation of H6. Thus, the engagement of β-CTT with H5 explains how isotype-specific differences in tubulin can be translated into functional changes at H6 and, consequently, into differences in dynein binding affinity. Importantly, this displacement of the MTBD H6 towards the α-tubulin was absent in the other isotypes (TUBB3, TUBB4A, TUBB5), where the β-CTT of the PF_2_ did not interact with the MTBD, and correspondingly, H6 remains disengaged from the MT surface, indicating that this effect is specific to engagement of the β-CTT with the MTBD. **(Table S7)**

The displacement of helix H6 in the TUBB2A, TUBB2B, and TUBB2C isotypes stabilizes MTBD binding to the MT and enhances binding affinity. This enhanced affinity provides a structural explanation for dynein’s increased processivity on MTs in the presence of the CTTs. Experimental studies have shown that enzymatic cleavage of tubulin CTTs significantly reduces the dynein binding to the MTs (by three-fold) and dynein’s run length (by four-fold), and indicated the role of E-hooks in mediating a weak interaction between the dynein and MTs (30). The structural explanation for these experimental observations is that the β-CTT enhances dynein motility by steering MTBD H5 and H6 helices into their high-affinity orientation, thereby strengthening MTBD-MT binding and increasing dynein binding and its run length. By cleaving the CTTs, the allosteric signal that engages helix H6 is lost, leading to a weaker MTBD-MT interaction and, consequently, a reduced run length.

Collectively, the findings demonstrate that the β-CTT of the neighboring PF can induce conformational changes in the MTBD, primarily by stabilizing helix H6 in conformations characteristic of the high-affinity state. These changes depend on the strength and duration of the β-CTT interaction with the MTBD, as indicated by the occupancy (i.e., percentage of the trajectory time during which an interaction is present) of the interactions between the α-tubulin and the MTBD H6. (**Table S7**)

In TUBB2B and TUBB2C isotypes, stable salt bridges and hydrogen bonds are formed between the β-CTT and the MTBD lasting for 150 to 200 ns after convergence, leading to an apparent displacement of MTBD H6 towards the MT, resulting in >50% occupancy (i.e., percentage of simulation time an interaction occurs) of MTBD H6-α-tubulin interactions. (**Figure S5B–C, Figure 13B and D, Figure S10B and D and Table S7**)

In contrast, in TUBB2A, the β-CTT formed only transient hydrogen bonding (lasting around 80 ns) with the MTBD, which resulted in a small displacement of H6 toward α-tubulin (as compared to the displacement of MTBD H6 in TUBB2B and TUBB2C tetramers) and a maximum of ∼30% occupancy (i.e., percentage of simulation time an interaction occurs) of α-tubulin interactions with the MTBD. (**Figure S5A, Figure S9B and D and Table S7**)

This conclusion is further supported by the average non-bonded interaction energy (electrostatic and van der Waals energy) calculated between the MTBD and the tubulin heterodimer on PF_1_. TUBB2B (−1382.33 kJ/mol) and TUBB2C (−1355.46 kJ/mol) displayed significantly more favorable energies than the TUBB2A isotype (−1138.76 kJ/mol). **(Figure S11)**

This difference in binding strength quantitatively supports the notion that strong (i.e., salt bridges and hydrogen bonds) and prolonged (for >150 ns) interactions with the β-CTT of the adjacent PF (e.g., PF_2_) enhance the binding affinity of the MTBD to the MT, promoting structural rearrangements characteristic of the high-affinity dynein-binding conformation.

Additionally, this investigation revealed a distinctively novel role of the lateral interactions between β-tubulin subunits of adjacent parallel filaments (PF_1_-PF_2_), which significantly influence the positioning and interaction of the β-CTT with the microtubule binding domain (MTBD) that was not previously reported. (**Figure 7**)

Differences in the β-tubulin core between isotypes altered the stability of the PF interface and indirectly modulated β-CTT-MTBD interactions. For example, TUBB2B and TUBB2A have identical β-CTT sequences, yet the β-CTT in TUBB2B formed persistent interactions with the MTBD (lasting over 150 ns; multiple salt bridges and hydrogen bonds). In comparison, in TUBB2A, it interacted only transiently (lasting ∼80 ns; hydrogen bonds and van der Waals interactions). (**Figure S2, Figure S5A–B, and Figure 5A–B**)

This divergence can be attributed to two amino acid differences (at positions 55 and 201) in the β-tubulin core between TUBB2A and TUBB2B isotypes, particularly at position 55, a residue located at the lateral interface. (**Table S5**)

In TUBB2B, this position is occupied by threonine (Thr55), which, due to its hydroxyl-containing side chain, acts as both a hydrogen bond donor (for Gln280, Arg282, and Gly360 of β-tubulin in PF_2_) and acceptor (for Gln280 of β-tubulin in PF_2_) and participates in three long-lasting hydrogen bonds at the lateral interface. (**Table S6**)

Conversely, in TUBB2A, this position is an alanine (Ala55), which lacks a polar side chain and forms only one long-lasting hydrogen bond with Gln280 of β-tubulin in PF_2_. (**Table S6**)

These findings suggest that enhanced lateral interactions in TUBB2B (2 salt bridges and 14 hydrogen bonds) stabilize the PF interface more effectively, restricting interdimer angles and positioning the β-CTT of the PF_2_ more favorably relative to the MTBD, thereby displacing the MTBD H6 effectively towards the MT. Thus, the strength of lateral interactions indirectly regulates β-CTT-MTBD binding by modulating PF geometry, thereby impacting dynein’s binding conformation and function.

## 4. Conclusions

This study investigates the structural roles of *Homo sapiens* tubulin isotypes, TUBB2A, TUBB2B, TUBB2C, TUBB3, TUBB4A, and TUBB5, in modulating dynein interactions with the MT, focusing on the interactions of their β-tubulins’ CTTs with the MTBD. The study identified the species-specific interactions between the β-CTT of PF_1_ and the β-tubulin core of the same PF in *Homo sapiens*, which differed from interactions observed in other species (*Sus scrofa* and *Bos taurus*), providing structural evidence that E-hooks can have a different effect on the MT-MTBD interactions even in closely related species due to the differences in the amino acid sequences at the C-terminal tails.

The analysis further demonstrated that the strength of lateral interactions between the β-tubulin subunits of the neighboring PF_1_ and PF_2_, based on the number of hydrogen bonds and salt bridges and non-bonded interaction energy, directly governs the degree of PF bending and interdimer angle fluctuations, significantly affecting the spatial positioning and orientation of the β-subunit of the PF_2_, particularly in relation to the MTBD bound on PF_1_. The results revealed that tubulin isotypes TUBB3, TUBB4A, and TUBB5 have relatively weak lateral interactions (№. hydrogen bonds ≤9) between the β-subunits of adjacent PFs (as compared to TUBB2A, TUBB2B, and TUBB2C isotypes), which led to rotation of the β-tubulin in PF_2_ around the MT longitudinal axis as indicated by more variable interdimer angles (ranging from 40° to 120°), causing the outward (i.e., away from the PF_1_) movement of the C-terminal helices (β-H15, H16, and H17) and β-CTT of PF_2_, thereby distancing the β-CTT from the MTBD. Conversely, isotypes (TUBB2A, TUBB2B, and TUBB2C) with stronger lateral interactions (№. hydrogen bonds ≥11 and №. salt bridges ≥1) maintained smaller interdimer angles (20°–80°) and structural stability, allowing sustained and functionally significant interactions between the β-CTT of PF_2_ and the MTBD.

Principal Component Analysis (PCA) and Dynamic Cross-Correlation (DCC) maps further clarified the allosteric coupling between the β-CTT and the MTBD. In TUBB2A, TUBB2B, and TUBB2C, we observed apparent anti-correlated motions between the MTBD and PF_2_, suggesting coordinated structural dynamics that facilitate interactions between the β-CTT and MTBD, consequently MTBD conformational switching.

Notably, the β-CTT interactions in TUBB2A, TUBB2B, and TUBB2C isotypes led to distinct rearrangements in key MTBD helices (H5 and H6), shifting the MTBD toward a conformation resembling the high-affinity state. These rearrangements include the reorientation of MTBD H5 by 10–40° at the YZ surface, the movement of MTBD H6 into proximity with α-tubulin, and the strengthening of the interactions between MTBD’s H6 and H16 and H17 helices of α-tubulin in PF_1_. These transitions are consistent with known structural determinants of high-affinity dynein binding reported through previous experimental studies.

Importantly, the differential ability of β-CTTs to induce these conformational changes was directly correlated with the duration and strength of their interactions with the MTBD, underlining the functional relevance of isotype-specific lateral bonding. TUBB2B and TUBB2C isotypes, which formed stable salt bridges and hydrogen bonds with MTBD (for ≥150 ns), induced the most pronounced conformational changes in the MTBD. In contrast, TUBB2A, with fewer (for ≤80 ns) and weaker interactions (only hydrogen bonds and van der Waals interactions), produced a slight reorientation of MTBD H6.

In summary, this work highlights the novel and unique role of β-tubulin isotype-specific lateral interactions, owing to the amino acids’ differences at the lateral interface in regulating dynein-MT dynamics’ by modulating PF geometry and enabling or restricting β-CTT-MTBD interactions, different tubulin isotypes can tune the conformational state of the MTBD and, by extension, dynein motor protein function, which has direct implications for the functioning of the dynein depending upon the tissues where these isotypes are expressed.

## Supplementary Information

**Figure S1:** Depiction of secondary structures of (A) αIA-tubulin (TUBA1A) and (B) βIII-tubulin (TUBB3) of 5JCO entry, and (C) Microtubule Binding Domain (MTBD) and the stalk domain of *Homo sapiens*. The secondary structures were determined through the ProS^2^Vi software package. The amino acids are shown as single-letter codes. The helices are represented in red, and the beta-strands are depicted in blue arrows.

**Figure S2:** Comparative sequence alignment of *Homo sapiens* β-tubulin wild type TUBB3 and isotypes TUBB2A, TUBB2B, TUBB2C (TUBB4B), TUBB4A, and TUBB5 performed using UniProt’s Clustal Omega algorithm. The amino acids are shown as single-letter codes. The asterisk (*) represents identical amino acids for all the sequences at that position. Colon (:) denotes a conservative mutation, period (.) represents a semi-conservative mutation, and blank space () indicates a non-conservative mutation.

**Figure S3**: Comparative sequence alignment of MTBD and stalk domain of the *Homo sapiens* (UniProt entry Q14204) and *Mus musculus* (UniProt entry Q9JHU4) performed using UniProt’s Clustal Omega algorithm. The amino acids are shown as single-letter codes. The asterisk (*) represents identical amino acids for all the sequences at that position. Colon (:) denotes a conservative mutation, period (.) represents a semi-conservative mutation, and blank space () indicates a non-conservative mutation.

**Figure S4**: Root mean square deviation (RMSD) analysis of different tubulin molecular systems (S1–S12). (A) TUBB2B α, β-tubulin heterodimer using GROMOS (S1, dark blue) and CHARMM (S2, orange) force fields, and TUBB3 heterodimer with GROMOS (S3, teal) force field (B) TUBB3 α, β-tubulin heterodimer with CHARMM (S4, red) and TUBB5 heterodimer with GROMOS (S5, dark green) and CHARMM (S6, light blue) force fields, (C) TUBB2A (S7, light green), TUBB2C (S9, dark brown), and TUBB4A (S11, purple) α, β-tubulin tetramer using GROMOS force field, and (D) TUBB2B (S8, light brown), TUBB3 (S10, pink), and TUBB5 (S12, olive green) α, β-tubulin tetramer with GROMOS force field. Solid lines indicate average trendlines generated by averaging every 500 frames.

**Figure S5**: Schematic representing the minimum distance between the β-CTT of the PF_2_ and the MTBD bound to the PF_1_ in (A) TUBB2A (S7), (B) TUBB2B (S8), (C) TUBB2C (S9), (D) TUBB3 (S10), (E) TUBB4A (S11), and (F) TUBB5 (S12) isotypes. The horizontal red dashed line represents the cut-off distance for the hydrogen bond (i.e., 0.35 nm), the cyan dashed line represents the cut-off value for the salt bridge interactions (i.e., 0.4 nm), and the horizontal dashed green line represents the cut-off (i.e., 1.4 nm) for long-range van der Waals interactions according to the GROMOS force field. The solid blue line depicts the 500-point moving average line.

**Figure S6**: Interdimer angle (θ) between the adjacent PF_1_ and PF_2_ of α, β-tubulin tetramers for (A) TUBB2A (S7), (B) TUBB2B (S8), (C) TUBB2C (S9), (D) TUBB3 (S10), (E) TUBB4A (S11) and (B) TUBB5 (S12) isotypes. The solid line represents the 500-point moving average line.

**Figure S7:** Projection of the backbone atom motions of (A–C) TUBB2A, (D–F) TUBB2B, and (G–I) TUBB2C isotypes in phase space along the first three principal components (PC1 vs. PC2, PC2 vs. PC3, and PC1 vs. PC3), colored from blue to red in order of time (from post-convergence to 1 μs).

**Figure S8:** (A) and (C) Electrostatic potential surfaces of *Homo sapiens* TUBA1A/TUBB2B tetramer and dynein (i.e., MTBD and stalk) from the PC1-derived conformation of the TUBB2B isotype. Red indicates negatively charged surfaces, while blue shows positively charged surfaces. (B) Negatively charged and aromatic amino acids (green sticks) in the β-H17 region and β-CTT of PF_2_ that generate the strong acidic potential in this region. (D) Positively charged amino acids on MTBD helices H2, H3, H5, and on the stalk’s CC1 and CC2 that areoriented towards the β-CTT of PF_2_. The electrostatic map is shown for TUBB2B, as the β-H17 and β-CTT region is similarly acidic across all isotypes, and the MTBD and stalk domains are identical in all isotypes.

**Figure S9:** Comparison of two distinct conformations of the TUBB2A tetramer from the first principal component analysis (PCA) (light blue and orange yellow) with the initial trajectory frame (α-tubulin: purple, β-tubulin: green, MTBD/stalk: gray). Panels (A) and (C) show the overall conformational differences of conformation 1 (light blue) and conformation 2 (orange yellow), respectively, while panels (B) and (D) present close-up views of the MT-MTBD interface in these two conformations with red and blue arrows showing the movement of MTBD helices H5 and H6, respectively.

**Figure S10:** Comparison of two distinct conformations of the TUBB2C tetramer from the first principal component analysis (PCA) (light blue and orange yellow) with the initial trajectory frame (α-tubulin: purple, β-tubulin: green, MTBD/stalk: gray). Panels (A) and (C) show the overall conformational differences of conformation 1 (light blue) and conformation 2 (orange yellow), respectively, while panels (B) and (D) present close-up views of the MT-MTBD interface in these two conformations with red and blue arrows showing the movement of MTBD helices H5 and H6, respectively.

**Figure S11**: Non-bonded (electrostatic and van der Waals energy) interaction energy between the α, β-tubulin heterodimer on the PF_1_ and the MTBD in the (A) TUBB2A, (B) TUBB2B, and (C) TUBB2C tetramer. The solid line represents the 500-point moving average line.

**Table S1**: Summary of the tubulin molecular systems (S1–S12) before subjecting to MD simulations, including the number of protein atoms, number of water molecules, number of sodium ions added, and the water box dimensions of each tubulin system.

**Table S2**: The energy minimization (EM) input (emtol and emstep) and output parameters (potential energy, maximum force, and normal of force) of different tubulin molecular systems (S1–S12) subjected to energy minimization using the steepest descent algorithm.

**Table S3**: Salt bridges and hydrogen bonds observed between the β-CTT of the PF_2_ and the H2–H3 loop (amino acids 3328–3334), H4–H5 loop (amino acids 3353–3360), and helix H5 (amino acids 3361–3371) of the MTBD bound to the PF_1_ in the TUBB2B tetramer (S8). The salt bridges were identified when the distance between oxygen atoms in the carboxyl group and nitrogen atoms in the amine group was less than the 4.0 Å cut-off at least once during the simulation trajectory. A hydrogen bond was defined using a donor-acceptor distance cut-off of 3.5 Å and a donor-hydrogen-acceptor angle cut-off of 30° (i.e., within 30° of linearity).

**Table S4**: Salt bridges and hydrogen bonds identified between the β-CTT of the PF_2_ and the H4–H5 loop (amino acids 3353–3360) and helix H5 (amino acids 3361–3371) of the MTBD bound to the PF_1_ in the TUBB2C tetramer (S9). The salt bridges were identified when the distance between oxygen atoms in the carboxyl group and nitrogen atoms in the amine group was less than the 4.0 Å cut-off at least once during the simulation trajectory. A hydrogen bond was defined using a donor-acceptor distance cut-off of 3.5 Å and a donor-hydrogen-acceptor angle cut-off of 30° (i.e., within 30° of linearity).

**Table S5**: Amino acids that differ in the β-tubulin core of the wild-type tubulin (TUBB3) and isotypes (TUBB2A, TUBB2B, TUBB2C, TUBB4A, and TUBB5), with corresponding secondary structure elements (H: α-helix, E: β-strand, and G: 3_10_-helix) where these amino acids are located.

**Table S6**: Long-lasting (occurring for >200 ns) salt bridges and hydrogen bonds found between the β-subunits of the PF_1_ and PF_2_ in wild-type and tubulin isotypes after the convergence. The salt bridges were identified when the distance between oxygen atoms in the carboxyl group and nitrogen atoms in the amine group was less than the 4.0 Å cut-off at least once during the simulation trajectory. A hydrogen bond was defined using a donor-acceptor distance cut-off of 3.5 Å and a donor-hydrogen-acceptor angle cut-off of 30° (i.e., within 30° of linearity).

**Table S7:** Hydrogen bonds found between the α-subunit of the PF_1_ and MTBD helix H6 (Tyr3379–Ala3385) and H5–H6 loop (Met3372–Asn3378) in TUBB2A, TUBB2B, and TUBB2C isotypes after the convergence. A hydrogen bond was defined using a donor-acceptor distance cut-off of 3.5 Å and a donor-hydrogen-acceptor angle cut-off of 30° (i.e., within 30° of linearity). The occupancy represents the percentage of trajectory time during which the hydrogen bond is present.

## Declaration of interests

The authors declare that they have no known competing financial interests or personal relationships that could have influenced the work reported in this paper.

## Acknowledgments

This study was funded by the Natural Sciences and Engineering Research Council of Canada (NSERC) Discovery Grant (No. **212654**) awarded to L.A.

The authors also thank the Digital Research Alliance of Canada and ACENET for the complimentary access to their High-Performance Computing (HPC) facilities and the technical support provided.

## Notes

### Competing Interest Statement

The authors have declared no competing interest.

